# Highly parallelized laboratory evolution of wine yeasts for enhanced metabolic phenotypes

**DOI:** 10.1101/2022.04.18.488345

**Authors:** Payam Ghiaci, Paula Jouhten, Nikolay Martyushenko, Helena Roca-Mesa, Jennifer Vázquez, Dimitrios Konstantinidis, Simon Stenberg, Sergej Andrejev, Kristina Grkovska, Albert Mas, Gemma Beltran, Eivind Almaas, Kiran R. Patil, Jonas Warringer

**Affiliations:** Department of Chemistry and Molecular Biology, University of Gothenburg, PO Box 462, 40530 Gothenburg, Sweden; European Molecular Biology Laboratory, Heidelberg, 69117, Germany; VTT Technical Research Centre of Finland Ltd; Espoo; 02044 VTT; Finland; Department of Biotechnology and Food Science, NTNU - Norwegian University of Science and Technology; Universitat Rovira i Virgili, Dept Bioquímica i Biotecnologia, Facultat d’Enologia, 43007 Tarragona, Spain; Centro Tecnológico del Vino—VITEC, Carretera de Porrera Km. 1, 43730 Falset, Spain; Medical research Council (MRC) Toxicology Unit, University of Cambridge, Cambridge, CB2 1QR, UK

**Keywords:** experimental evolution, genotype-phenotype relations, fermentation, metabolism, high-throughput, evolutionary engineering

## Abstract

Adaptive Laboratory Evolution (ALE) of microbes can improve the efficiency of sustainable industrial processes important to the global economy, but chance and genetic background effects often lead to suboptimal outcomes. Here we report an ALE platform to circumvent these flaws through parallelized clonal evolution at an unprecedented scale. Using this platform, we clonally evolved 10^^4^ yeast populations in parallel from many strains for eight desired wine production traits. Expansions of both ALE replicates and lineage numbers broadened the evolutionary search spectrum and increased the chances of evolving improved wine yeasts unencumbered by unwanted side effects. ALE gains often coincided with distinct aneuploidies and the emergence of semi-predictable side effects that were characteristic of each selection niche. Many high performing ALE strains retained their desired traits upon transfer to industrial conditions and produced high quality wine. Overall, our ALE platform brings evolutionary engineering into the realm of high throughput science and opens opportunities for rapidly optimizing microbes for use in many industrial sectors which otherwise could take many years to accomplish.

## INTRODUCTION

Microbial processes play important roles in the global economy with the production of fermented food and drinks alone accounting for trillions of dollars in turn-over^1–3^. Beyond fermented food and drinks, microbial fermentation is central to many production processes such as enzymes, anti- and probiotics, fine chemicals, and biofuels. However, the naturally occurring microbes seldom operate at efficiencies required for an economically viable production, leading to intense efforts to improve their properties^4–6^. Rational genetic engineering has helped in addressing some microbial shortcomings^7–9^, but generates genetically modified organisms (GMO) unsuitable for food, feed, or beverage production. Further, successful genetic engineering requires a deep understanding of the genotype-phenotype map^10–13^, which is often unknown, especially in the cases of complex, multi-genic traits. Mathematical modelling of cellular states and fluxes is equally challenging due to the interconnected nature of metabolic and regulatory processes^14^.Manifestation of both the desired industrial trait and unwanted side effects therefore often depend both on the genetic background and the environment^15–17^, and defy prediction.

Adaptive Laboratory Evolution (ALE) offers an attractive alternative for microbial improvement because it is unburdened by the need to understand the genotype-phenotype relation on a single gene level^18, 19^. As such, it has been successfully used for e.g. microbial thermotolerance^20^, methylotrophy^21^, carotenoids production^22^, and alcohol tolerance^23, 24^. Yet, ALE lineages often fail to evolve the desired traits, or end up carrying unwanted side effects, because etiology of the desired traits commonly involve neutral, costly, inaccessible, or highly pleiotropic mutations^25^ (Figure 1A). Moreover, chance influences both the birth and early fate of mutations, delaying the establishment of beneficial variants in ALE populations and allowing their neutral or weakly deleterious cousins to become common^2627^ (Figure 1B). The power of numbers could unshackle ALE from the constraints of chance, but achieving a sufficiently high ALE throughput without compromising error rates has proven challenging^27–31^. We therefore here developed an ALE platform capable of independently evolving 10^4^ microbial populations, and we demonstrate its utility by selecting for eight desired wine production traits while measuring adaptation, and its side effects, with high accuracy. The massive expansion of ALE replicate populations and parental lineages dramatically improved the chances of evolving well-adapted lineages unencumbered by slow growth side effects and many high performing ALE lineages retained the desired traits when transferred to industry-like conditions.

**Figure 1.**
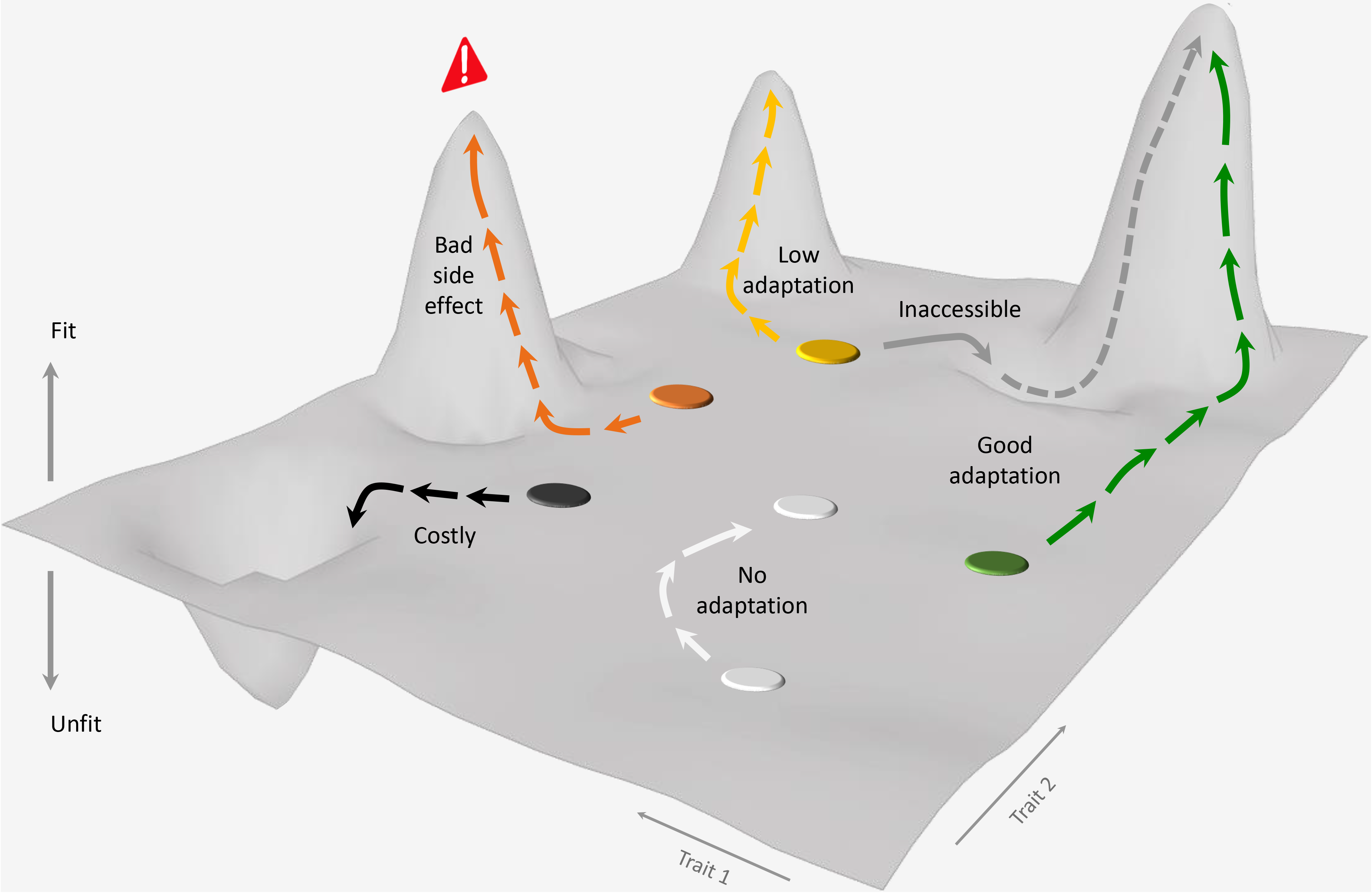
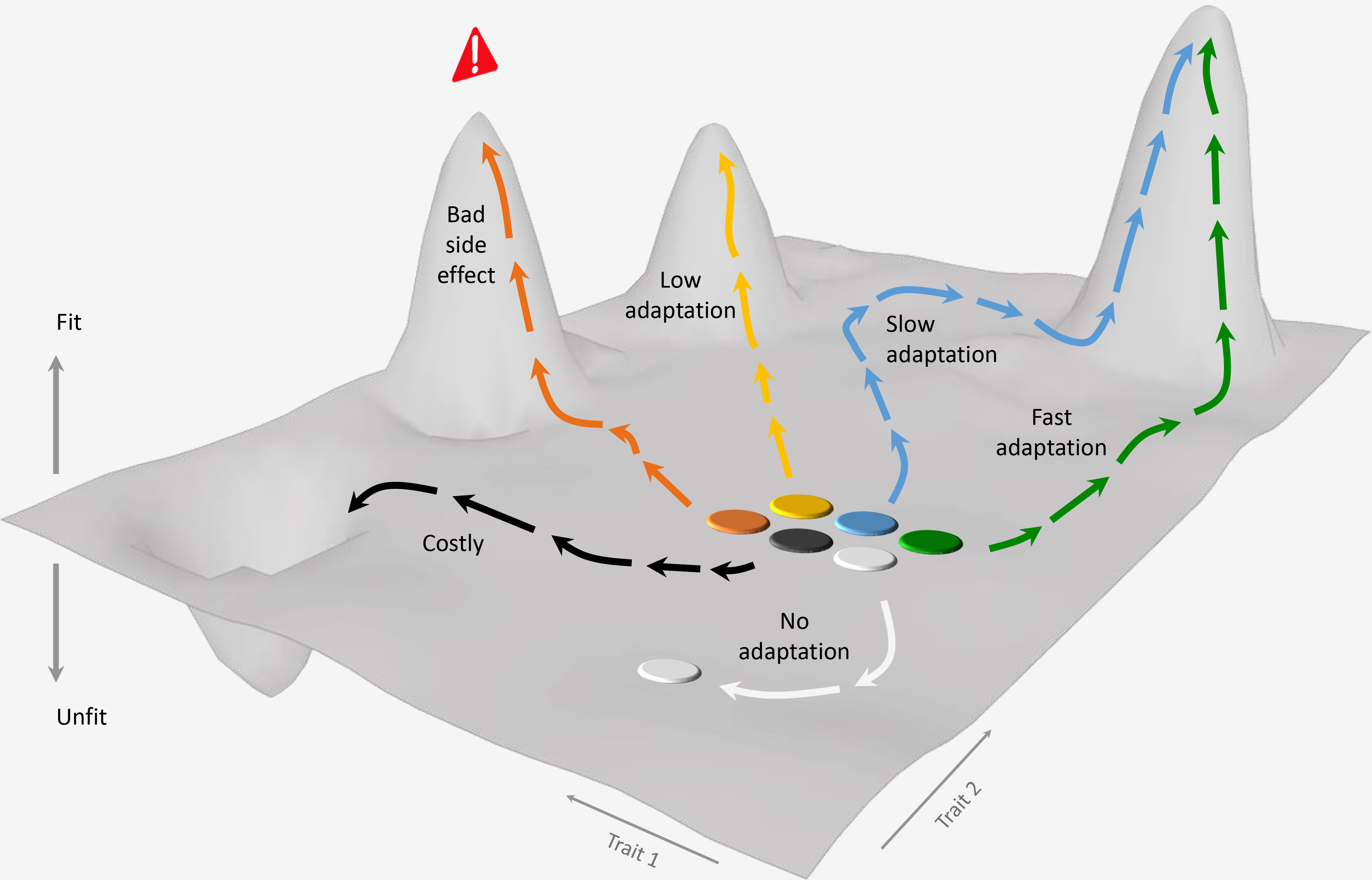
ALE challenges overcome by parallelization. **(A)** Mutations underlying desired traits may be neutral (blue), costly (black), inaccessible (grey), low adaptive (yellow), or associated with side effects (orange) in a lineage. Parallelizing ALE across many lineages can increase the chances of some being capable of evolving a desired trait that is unburdened by side effects (green). **(B)** Chance affects new mutations in a population, leading to slow (blue), or low yield (yellow), adaptation, and a side effect burden (orange). Parallelizing ALE across many replicated populations of a lineage can increase the chances of some quickly adapting to a high adaptation yield without becoming burdened by unwanted side effects (green).

## MATERIALS AND METHODS

### Strains and growth medium

We obtained 15 commercial wine yeasts marketed by Lallemand Inc. (Canada) and 33 pre-commercial wine yeasts. We isolated the latter from grapes or vineyard soil in the Priorat wine-making region in Catalonia and identified them as *S. cerevisiae* using restriction fragment length polymorphisms. Strains are listed in Table S1. Strains were stored long term at −80 C in 20% (v/v) glycerol. Experiments in Figures 2-4 and S2-6 were performed in synthetic grape must medium^32^. Vitamins (100x; pH adjusted to 3.3 with NaOH), amino acids (10x; pH adjusted to 3.3 with NaOH, buffered with 2 % (w/v) Na_2_CO_3_), oligo-elements (1000x) and anaerobiosis factors (1000x; ergosterol, oleic acid, Tween 80 and ethanol) were stored as separate, sterile filtered stock solutions at 4 C (amino acids <2 weeks; vitamins at −20 C). To prepare the synthetic grape must, glucose (100 g/L), fructose (100 g/L), citric acid (5 g/L), malic acid (0.5 g/L), tartaric acid (3 g/L), KH_2_PO4 (0.75 g/L), K_2_SO4 (0.5 g/L), MgSO4.7H_2_0 (0.25 g/L), CaCl_2_.2H_2_O (0.155 g/L), NaCl (0.2 g/L) and NH4Cl (0.153 g/L) were dissolved in H_2_O, autoclaved, and pH were set to 3.3 with NaOH. Stock solutions were added. For solid medium experiments, a separate solution of the gelifying agent gelrite (gellan gum), which have better retention of water, less phenolic contamination and better light transmission properties than the classical agent agar^33–36^, and CaCl_2_.2H_2_O (initiates gelification) were then prepared, adjusted to pH=3.2 (NaOH), autoclaved, and added to the medium (final concentrations: gelrite=8 g/L, CaCl_2_.2H_2_O=1.155 g/L). The volume was set to 1 L (w. sterile H_2_O). For solid medium, the medium was stirred on a heater (at 50 C) and 50 mL was poured into the same lower left corner of each Plus plate (Singer, UK). Plates were dried overnight and used, with no additional storage. For particular experiments, variations to the basic synthetic grape must were made, as described in Table S2 and S3. Scale-up experiments (5 L and 80 L) were performed in White Grenache (GR) grape must recently harvested in DO Terra Alta (Spain).

**Figure 2.**
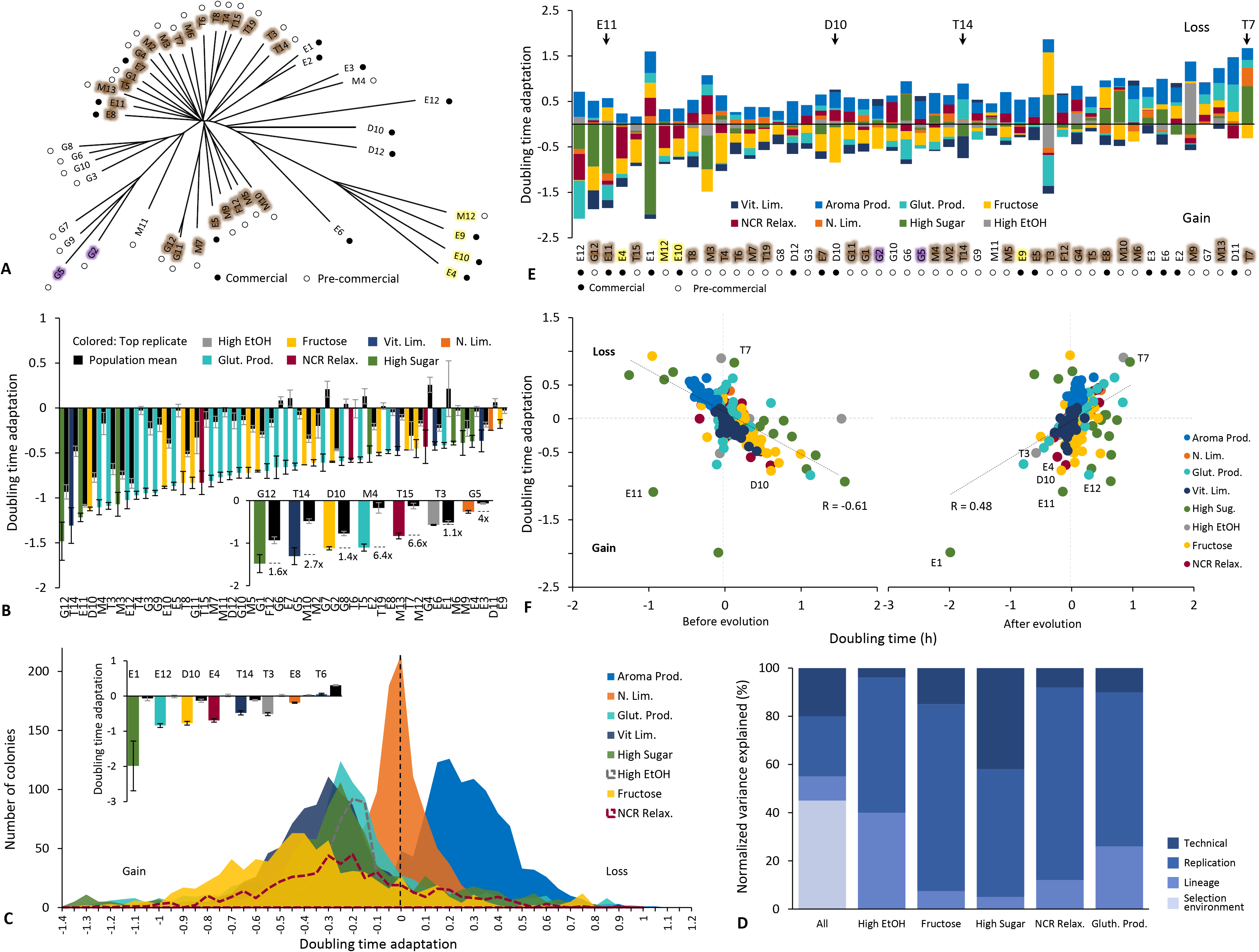
Wine yeast adaptation under highly parallelized ALE. We parallelized ALE across many (*n*=24) replicated populations of 48 (Table S1) commercial and pre-commercial wine yeasts as replicated colony populations on each of eight synthetic grape must conditions designed to select for traits desired by the wine industry (Table S2 and S3). The ALE design for the 9216 populations is shown in Figure S2. **(A)** Phylogeny (neighbor joining tree) of 47 commercial (●) and pre-commercial (○) euploid diploid wine yeasts, based on 42,599 single nucleotide polymorphisms called against the Wine/European strain EC1118. Coloured fields = sub-clades (see Figure S1). Bar = SNP distance. **(B-E)** We counted cells in growing populations to generate high density growth curves and estimated adaptation as the log(2) change in the cell doubling time from the respective founder strain to each ALE endpoint. **(B)** Adaptation (mean, *n=4*) of the most adapted replicated wine yeast lineage (left bar, indicated by name) as compared to the average lineage (*n=24*) across selection regimes. *Inset:* adaptation (mean, *n=4*) of the top replicate of the most adapted wine yeast lineage (left bar, indicated by name) as compared to the average lineage (*n=24*) in each selection ergim. FDR: q=0.05. Error bars: SEM. **(C)** Histogram of the adaptation of all ALE populations, in each selection environment (colour). Extinct ALE populations are not included. *Inset:* Mean (*n=24*) adaptation of the wine strain with the greatest adaptation (left bar, indicated by name) as compared to the average wine strain (*n=48*), in each selection environment. Error bars: SEM. **(D)** Percentage of the variance in adaptation that is explained by differences between: ALE selection environments, founder strains, replicated ALE populations, and technical replicates. **(E)** Stacked bar plot of the mean (*n*=24; extinct populations excluded) adaptation for each founder wine strain, in each ALE selection environment. ALE selection environments with positive adaptation (a cell doubling time decreases) are shown as negative sums, environments with negative adaptation (a cell doubling time increase) are shown as positive sums. Zero: no adaptation. Text colour: population (see Figure S1). Arrows: lineages indicated in text. **(F)** Comparing mean (*n*=24) adaptation, for each founder wine yeast in each ALE selection environment, to the mean relative doubling time before (left panel, founders) and after (right panel, *t_30_*) ALE selection. Colour=selection environment. Broken line = linear regression. Pearson’s *r* and lineages mentioned in text are indicated.

**Figure 3.**
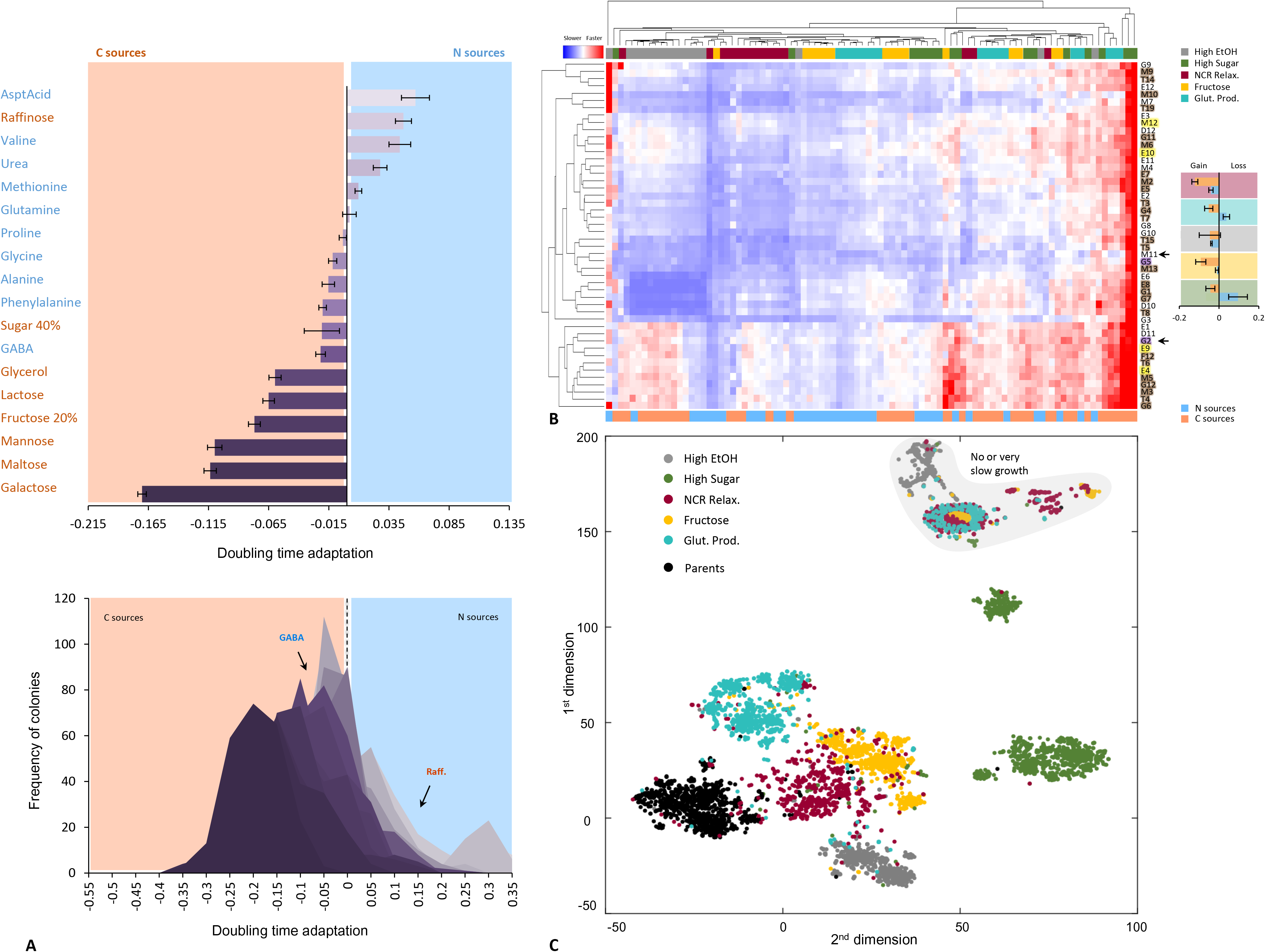
Side effects in ALE populations. We cultivated start and endpoints of ALE populations in eight selection environments in 18 non-selection carbon and nitrogen limited niches. We estimated ALE side effects as the log_2_ change in doubling time between the start and endpoint state, in these niches. **(A)** ALE side effects, averaged across all ALE replicates, lineages and selection regimes (*n*=5760). Faster growth side effect = negative numbers. Orange text: carbon limited niches, blue text: nitrogen limited niches. Error bars: SEM (across selection regimes, *n*=5) **(B)** *Central heat map*: ALE side effects evolved by each lineage in each selection regime. Each column represents one type of side effect (growth in a carbon or nitrogen limited niche), evolved under one selection regime. Each row represents one lineage (mean of *n=24* populations), with names coloured by clade (see Figure S1). Arrows indicate M11 and G2. Red = faster growth side effect, blue = slower growth side effect. *Upper panel:* Hierarchical clustering of side effect niches. Top colour panel: selection regime. Note that sets of side effects are grouped by selection regime. Bottom color panel: carbon (orange) and nitrogen (blue) limited niches. *Left panel:* Hierarchical clustering of lineages based on similarity across side-effect-environments combinations. **(C)** t-Distributed Stochastic Neighbor Embedding (t-SNE) clustering reducing the side effect variation to two dimensions. Each dot represents one ALE population, of one lineage, in one selection regime - or one founder strain. Colour = selection regime, as in (B). Note that side effects cluster by selection regime, representing common syndromes.

**Figure 4.**
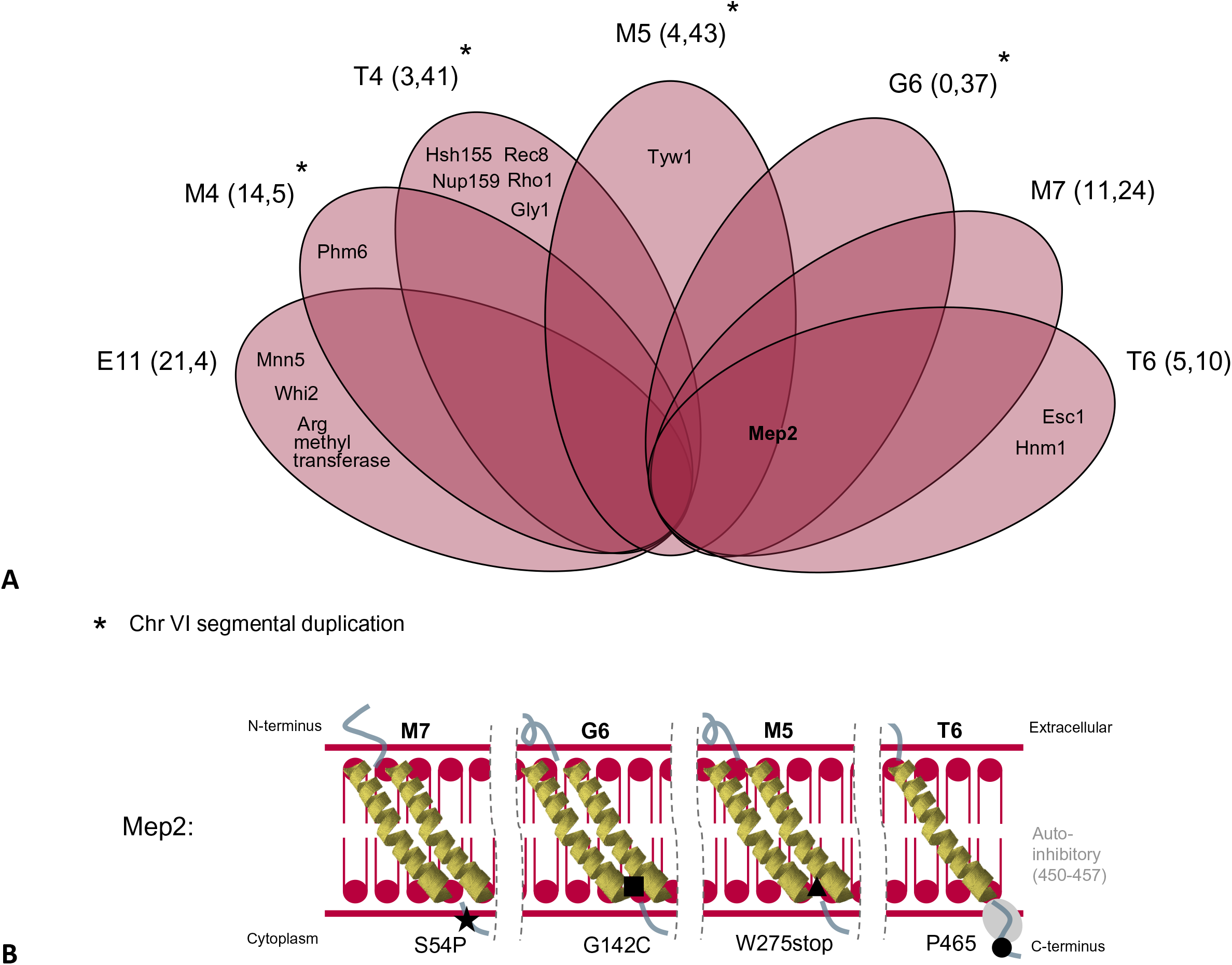
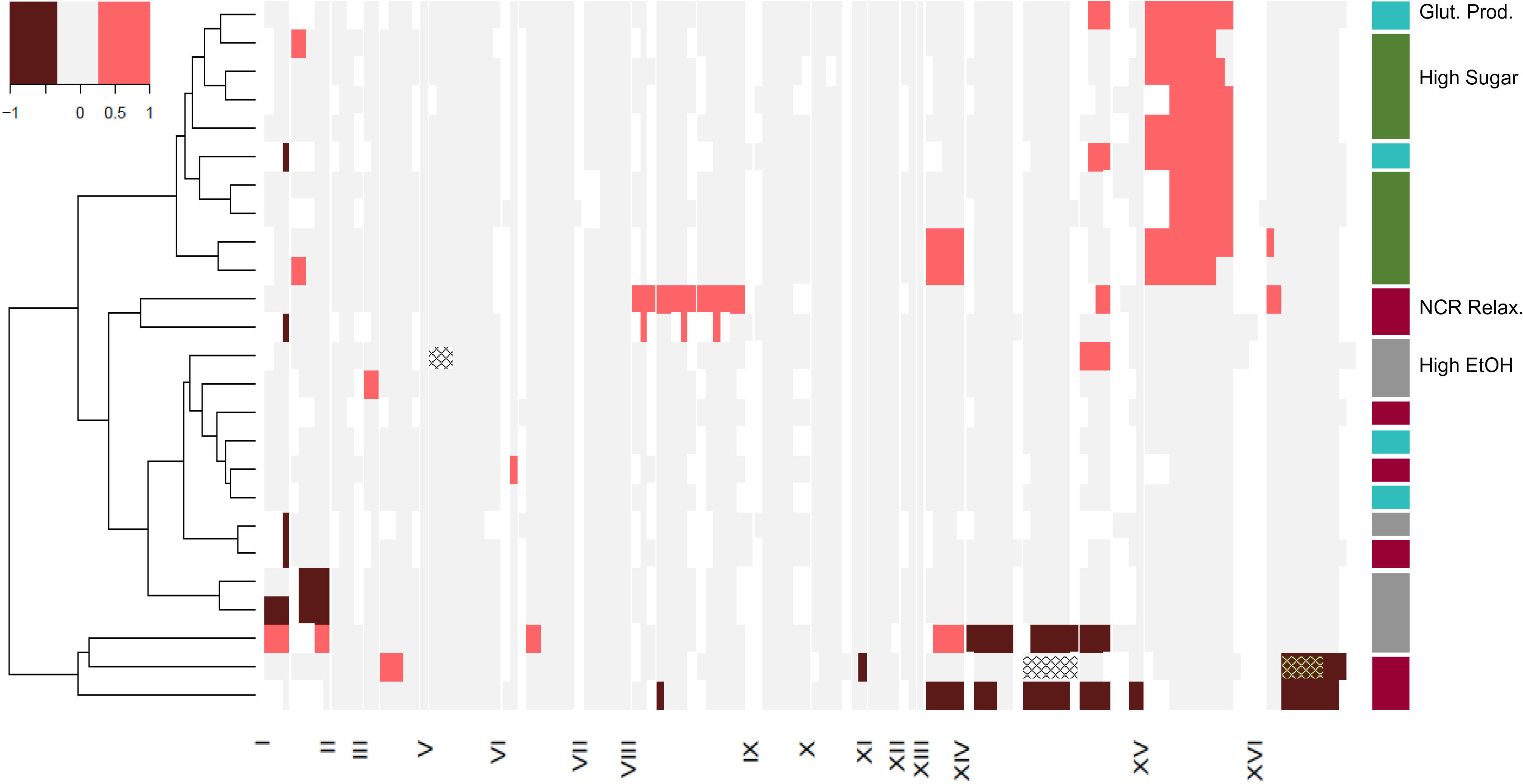
*MEP2* and chromosomal mutations drive ALE adaptations. **(A)** Venn diagram depicting proteins affected by non-synonymous mutations in seven sequenced populations ALE adapted for NCR relaxation. Names of populations (outside) and genes (inside) are indicated. Mep2 (bold) were mutated in four populations. **(B)** Schematic view of the high affinity ammonium and methylammonium permease Mep2. The seven transmembrane regions (yellow), the autoinhibitory domain (green) of the cytoplasmic tail and the amino acid mutations driving adaptation for NCR relaxation, and population names (top) are all indicated. **(C)** ALE wine yeast populations often acquired chromosome or chromosome segment copy number variations (CNV; colour) or loss of heterozygosity. Changes corresponding to 20% larger, or smaller, read coverage than expected are shown for 26 sequenced ALE populations. White vertical lines indicate ends of contigs.

### Adaptive Laboratory Evolution (ALE)

Each of the 48 strains to be used for ALE was stored as two separate populations, both clonally expanded from the same single cell, in 20 % (v/v) glycerol in 96 well format. The 96 frozen stocks were thawed, re-pinned 12x onto solid, synthetic grape must medium (Figure S2) using long-pin pads (Singer) to generate 1152 colonies in a 1536 array and cultivated for 72 h. This pre-culture was repeated once to further standardize the physiological states of populations. We maintained one in every four positions empty to continuously survey cross-contamination and to allow inclusion of fixed controls in the growth measurement stage. Standardized pre-cultures were re-pinned (1536 short pin transfer) onto eight selection environments to generate in total 9216 ALE yeast populations. The eight selection media were synthetic grape must with: (i) 20% (w/v) fructose as the sole sugar, together with 2 g/L 2-deoxy glucose (fructose utilization), (ii) 10% of the regular amino acid concentration, i.e. 10 mg N/L (nitrogen starvation), (iii) glycine, glutamine and cysteine as the sole nitrogen sources, together with 1.5 mM diamide (glutathione production), (iv) arginine and proline as the sole nitrogen sources together with 1% (w/v) methyl amine (Nitrogen Catabolite Repression (NCR) relaxation), (v) 35% (w/v) sugar with equal proportion of fructose and glucose (high sugar tolerance), (vi) 1.3% (v/v) 1-butanol (ethanol tolerance), (vii) valine, iso-leucine, and phenylalanine as the sole nitrogen sources (aroma production), and (viii) 1 % of the regular vitamin concentration (vitamin starvation). Synthetic grape must was used as pre-culture for the selection media (i), (v), and (vi), while synthetic grape must with low nitrogen (N) content (30 mg N/L) was the pre-culture for the rest of the selection media. We passed populations through a batch-to-batch selection regime of 30 consecutive growth cycles. In each cycle, we clonally expanded populations for 72 h from around *n=10^5^* cells, subsampled the mostly stationary phase populations using 1536 short pin transfers, and deposited samples (*n=10^5^* cells) on fresh selection medium. Each growth cycle corresponded to a mean of 6.6 population doublings for a total of ∼200 doublings (corresponding to cell generations, assuming no cell death). No invasions of presumed empty colony positions were observed, and a RFLP analysis of variation at the delta element locus of 65 fast-adapting endpoint populations showed that nearly all (n=56) retained the parent strain RFLP pattern (an unequivocal call could not be made for the remaining nine). Thus, while we cannot completely exclude cross-contamination between neighboring colony positions during the ALE, it should be rare.

The ALE procedure was standardized to minimize bias as follows. Solid media was cast in polystyrene plates from a single batch (Singer Instruments, SBS-format PlusPlates). Each plate was cast on top of the same perfectly level surface with precisely 50 mL of synthetic grape must medium at 50 C. Medium was poured in the same lower left corner on all plates. All plate preparations were performed in the same environmentally controlled space (room temperature 23 C). Pinning transfers were performed using 1536 short pin pads from the same production batch (Singer Instruments, UK), and a Singer RoToR HDA robot (Singer Instruments, UK). All transfers were performed using a single robot, standing in the same environmentally controlled space throughout the experimental series and a single production batch of Plus plates and pin pads. The robotic workspace was sterilized by prolonged UV exposure before each use and routinely cleaned with ethanol. All evolution plates were kept in the same temperature controlled (30 C) thermostatic cabinet. We stored a frozen fossil record of populations in glycerol in 96-well plates, by pinning from agar to liquid medium using the Singer RoToR HDA robot. After 3 days of incubation at 30 C, we added glycerol to a final concentration of 20% (v/v) and samples were stored at −80 C.

### Counting cells in growing founder and ALE endpoint populations

To monitor population size expansion of founder (*t*=0 growth cycles) and end-point (*t*=30 growth cycles) populations, we used a high-resolution microbial growth phenotyping platform. First, we measured the growth of founder populations in 70 environments (Figure S3) at high replication (*n=24*) by thawing of frozen stocks, pre-cultivation (2x), subsampling and deposition on fresh plates, as described above. Second, we thawed frozen stocks of founder and end-point populations, pre-cultivated these (2x) in parallel, subsampled pre-cultures deposited subsamples on each of 18 different environmental plates and measured their doubling time. Experimental populations in the two set-ups were handled and analyzed identically. Using a custom-made RoToR pinning program, we deposited 384 control samples, subsampled from a 384 pre-culture array of genetically identical control colonies (strain E9), in the 384 interleaved empty positions on each target plate. We recorded population size at 20 min intervals for 3 days using high-quality desktop scanners (Epson Perfection V_800_ PHOTO scanners, Epson Corporation, UK) connected via USB to a standard desktop computer. We performed transmissive scanning at 600 dpi using 8-bit grey-scale. Plates were fixed in place by custom-made acrylic glass fixtures. Pixel intensities were normalized and standardized across instruments using transmissive scale calibration targets (Kodak Professional Q-60 Color Input Target, Kodak Company, USA) which were fixed to each fixture. Pixel intensities were estimated and summed across each colony, subtracting the local background, and converted into cell counts by calibration to independent cell number estimates obtained using spectrometer and flow cytometry. Raw measurements of population size were smoothed, and the steepest slope in each growth curve was converted into a population size doubling time. Noisy growth curves were flagged, visually inspected for artifacts and 0.3% of doubling time estimates were rejected as potentially incorrect.Doubling times were log_2_ transformed and normalized to those estimated for each position using the 384 spatial controls on each plate, thereby producing log_2_ doubling time ratios that account for systematic variations in doubling times across and between plates.

### Sequencing and sequence analysis

Total genomic DNA was extracted from founder strains grown in YPD using phenol-chloroform based extraction and from evolved populations grown in YPD using Epicentre MasterPure Yeast DNA Purification Kit. The DNA quality was evaluated with electrophoresis in a 1% (w/v) agarose gel and DNA concentrations were evaluated using Qubit (Thermo Fisher Scientific, USA). An equal amount of DNA from each sample was used for library preparation with the NEBNext DNA Ultra2 Library Preparation Kit (New England Biolabs). The library preparation was performed on an automated liquid handling system (Hamilton Robotics) and the quality of the library was tested on a 2100 BioAnalyzer (Agilent Technologies). Paired-end Illumina short read sequencing was performed at the Genomics Core Facility (EMBL Heidelberg) on HiSeq 2000 and HiSeq2500 platforms (Illumina, San Diego, USA) for 150 bp (average insert size: 245 bp) and 250 bp (average insert size: 616 bp) reads, respectively. The data have been deposited in the European Nucleotide Archive (ENA) at EMBL-EBI under accession number PRJEB41108 (https://www.ebi.ac.uk/ena/browser/view/PRJEB41108).

The quality of the sequencing reads was controlled using Fastqc v. 0.11.4 (https://www.bioinformatics.babraham.ac.uk/projects/fastqc/; Andrews, 2010). The reads were trimmed by removing adapters and filtering low quality reads using cutadapt v. 1.10^38^ (https://pypi.org/project/cutadapt/). The trimmed reads were aligned to the *S. cerevisiae* EC1118 wine yeast regenome assembly^3942^ with the Burrows-Wheeler Aligner v. 0.7.12^40, 41^ using default parameters. Picard Tools v. 1.129 (https://broadinstitute.github.io/picard/) were used to process (read groups addition, sorting, reordering, and indexing) the alignments and mark duplicate reads.

Single nucleotide polymorphism (SNP), and insertion-deletion (indel) variant calling of founder samples (excluding strain D11 with low quality sequencing sample) was performed with GATK4 v. 4.1.0.0 *HaplotypeCaller* in GVCF model using the *S. cerevisiae* EC1118^39^ as the reference with DISCOVERY genotyping mode, ploidy 2, and minimum base quality score 20. The individual GVCF files were then combined using *CombineGVCFs*, and jointly genotyped using *GenotypeGVCFs* with ploidy 2 and standard minimum confidence of calling 20. Using *SelectVariants* and *VariantFiltration* tools the called SNPs and indels were filtered separately. SNPs were filtered with QD < 2.0, FS > 60.0, MQ < 40.0, MQRankSum < −12.5, ReadPosRankSum < −8.0, GQ < 30, DP < 5. Indels were filtered with QD < 2.0, FS > 200.0, MQ < 40.0, ReadPosRankSum < −20.0, GQ < 30, DP < 5. Then, the SNP sites were further filtered using vcftools v. 0.1.14 for sites which miss more than 50% of the data (max-missing, 0.5), for other than biallelic sites (max-alleles, 2; min-alleles, 2), for sites with minor allele frequency less than 0.05 (maf, 0.05), and for sites not in Hardy-Weinberg equilibrium (hwe, 0.00001). To estimate average nucleotide diversity among the parent strains vcftools v. 0.1.14 was used. A neighbour-joining tree of the founder strains was created from the filtered set of 42599 segregating sites using R v. 4.0.3^42^ and packages ape v. 5.4.1 (https://github.com/emmanuelparadis/ape) and SNPrelate v. 1.24.0 (http://github.com/zhengxwen/SNPRelate). First, individual dissimilarities (derived from relative coancestry) were estimated for each pair of individuals with the snpgdsDiss function. The individual dissimilarities were then used as distances for the bionj function.Model-based Bayesian algorithm fastSTRUCTURE v. 1.0^43^ was used to detect and quantify admixture in the 47 parent strain genomes (excluding strain D11 with low quality sequencing sample). fastSTRUCTURE was run on the filtered set of 42599 segregating sites, varying the number of founder populations (K) between 1 and 10 using the simple prior implemented in fastSTRUCTURE. K=4 was found to be optimal, i.e. scoring the highest marginal likelihood.

Single nucleotide variant iant (SNV), and small insertions-deletions (indel) in ALE populations were called using GATK4 Mutect2 v. 4.1.0.0^44^, default parameters and a panel-of-normals approach. We created the high coverage sequence panel using 47 parent strains (excluding D11) and the *CreateSomaticPanelOfNormals* tool. We called variants in each ALE endpoint relative to the panel using *FilterMutectCalls* and default parameters, including a tumor-lod of 5.3. Copy number variants in ALE populations (CNV) were called analysis using ATK4 v. 4.1.0.0 tools. Read for 1000 bp intervals were counted with *CollectReadCounts and* read counts were denoised and matched to that of its corresponding parent strain sample using *DenoiseReadCounts*. The allelic counts were collected using *CollectAllelicCounts* and combined with the binned read count ratios for modelling the CNV segments using *ModelSegments* with number-of-change-points-penalty-factor of one (1) or ten (10) for HiSeq 2000 and HiSeq2500 samples respectively. CNVs were called using *CallCopyRatioSegments*. Called CNV segment copy ratios were further re-centered to the corresponding sample medians. The sample H2, with a high noise copy ratio comparing to the corresponding parent sample and whose contigs were smaller than 10 kb was excluded. Loss-of-heterozygosity in ALE populations was called as baf zero/one segments across sites for which the corresponding parent strain was called as hetorozygotic (baf∼0.5).

### Fermentation validation of ALE endpoints

Selected ALE endpoint populations were validated at lab scale by fermenting them and their respective parent strains in a synthetic grape must (pH 3.3) as described in Beltran et al.^32^, with the nitrogen content varying between fermentations. The control synthetic grape must contained 200 g/L of sugar (100 g/L glucose and 100 g/L fructose) and 300 mg of yeast assimilable nitrogen/L (120 mg/L inorganic and 180 mg/L organic nitrogen).

All fermentations were performed as biological triplicates at 22 C with agitation (120 rpm) in laboratory-scale fermenters: 250 mL flasks containing 200 mL of medium and capped with closures that enabled carbon dioxide to escape and samples to be removed. Fermentations were performed in semi-anaerobic conditions. The initial yeast inoculum consisted of 2 × 10^6^ cells/mL taken from YPD stationary phase cultures (48 h). Sugar consumption was monitored throughout the fermentation process by measuring decrease in fermentation medium density using a densitometer (Densito 30PX, Mettler Toledo, Switzerland). In the later stages of the fermentation, when the sugar contribution to medium density is limited, sugar consumption (glucose and fructose) was assayed using enzymatic kits (Roche Applied Science, Germany). Fermentation was considered to be completed when residual sugars were below 2 g/L. Yeast cell counts were determined by measuring optical density at 600 nm.

Yeasts cells were harvested at different time points for measurement of the nitrogen, sugar or total glutathione content. Cell pellets were transferred to Eppendorf tubes, frozen immediately in liquid nitrogen and kept at −80 C until they were analysed. The supernatant was stored at −20 C for later analysis of their remaining nitrogen content.

### Semi-industrial validations of ALE endpoints

Semi-industrial fermentations were conducted in 100 L stainless steel tanks filled with 80 L of White Grenache (GR) grape must from DO Terra Alta (Spain), as well as in 5 L fermenters, filled with 4 L of the same grape must. The musts had a density around 1100 g/L, pH=3.31, and an initial yeast assimilable nitrogen content of 163.8 mg/L. 80 L fermentations (*n*=1) and 5 L fermentations (*n*=3) were performed using single colonies isolated from selected ALE endpoint populations and their corresponding founders, as listed in Table S4. From each tank and fermenter, daily samples were taken to monitor sugar concentration by measuring must density using an electronic densitometer (Mettler-Toledo S.A.E., Barcelona, Spain), as well as yeast growth (by counting of colony forming units). The final wines were stabilized for 30 days at 4 C, 30 ppm of sulfur dioxide was added as potassium metabisulfite, and the product was bottled and stored for 2 months until the sensory evaluation took place. At the middle and end point of the fermentation, *S. cerevisiae* isolates were typified by the analysis of inter-delta regions, as described by Legras and Karst (2003)^45^ using primers Δ12 and Δ21.

### Nitrogen content analysis

Selected ALE endpoint populations and their corresponding parents (see Figure S9B) were cultivated as 80 mL cultures. Free amino acids and ammonia were analysed using DEEMM derivatizations as in Gomez-Alonso et al.^46^ and with the Agilent 1100 Series high-performance liquid chromatography (HPLC) (AgilentTechnologies, Germany). Separation was performed in an ACE HPLC column (C18-HL) particle size 5 µm (250 mm x 4.6 mm) thermostatized at 20 C. Several dilutions of each sample were analysed and averaged. The concentration of each compound was calculated using an internal standard, Agilent ChemStation (Agilent Technologies) and expressed as mg N/L.

### Glutathione measurement

Total glutathione levels (intracellularly and extracellularly) were established in stationary phase, in 18 ALE endpoint populations and their founder strains (Table S5). Cells were pre-cultivated on 50 mL of glutathione evolution medium (see above) at 28 C with orbital shaking (120 rpm) and after 48 h growing, cells were inoculated in 45 mL of the same medium at initial OD_600_=0.2 (*n=2*). Yeast growth and media density was evaluated continuously by spectrometry and electronic densitometry and glutathione was measured after one week, with all cultures being well into the stationary phase.

For glutathione extraction, the method described by Vázquez et al., 2017 was used^47^. Briefly, a cell density of 1×10^8^ cells was centrifuged at 10,000 rpm to obtain a packed cell pellet and the supernatant. Pellets were weighed and three volumes of 5% (v/v) 5-sulfosalicylic acid (SSA) were added and vortexed. The cell suspensions were then frozen and thawed three times (using liquid nitrogen to freeze and a 37 C bath to thaw), incubated for 5 min at 4 C and centrifuged at 800 g for 10 min at 4 C. The measurement of total glutathione (tGSH) was determined (in pellets and supernatants) using the kinetic glutathione assay kit (Sigma-Aldrich, Spain) in which catalytic amounts (nmoles) of reduced glutathione (GSH) cause a continuous reduction of 5,5’-diothiobis-2-nitrobenzoic acid (DTNB) to 5-thio-2notrobenzoic acid (TNB), and the oxidized glutathione (GSSG) formed is recycled by glutathione reductase and NADPH. Final TNB formed (equivalent to tGSH) was measured spectrophotometrically at 412 nm. Other pellets (1×10^8^ cells) were previously dried at 28 C for 48 h and weighed. Thus, the results are expressed as nmoles tGSH/mg dry weight.

## Results

### Parallelizing ALE for high throughput

We developed a highly parallelized ALE platform and evaluated its applicability by evolving 48 wine yeasts for eight desired wine yeast traits. The selected traits include growth on less-preferred nutrients (fructose and arginine, proline), growth on eccentric concentrations (high sugar, high ethanol, low vitamin and low nitrogen) and production of metabolites (glutathione and aromatic compounds) towards a more efficient and better quality wine production for which we designed specific environments (Table S2 and S3). We used 15 commercial wine yeasts (Lallemand Inc. Canada) and 33 pre-commercial lineages isolated from cellars across the Priorat wine district in Spain as genetic starting points (Table S1) and sequenced their genomes, depositing data in the European Nucleotide Archive (Methods). Commercial and pre-commercial lineages all shared recent ancestry (nucleotide diversity, π=0.000018 to 0.011) within the Wine/European clade, but dispersed into four sub-clades with only limited admixture^48, 49^ (Figures 2A, S1A). Uniform allele frequency ratios of ∼1:1 showed all to be euploid diploids with only a few small and scattered segmental aneuploidies. We repeatedly (*n*=24) expanded all 48 lineages from clones to generate 1152 founder populations and cultivated these as colonies on solid plates based on each of the eight synthetic grape must media^32^ designed to select for the different desired traits (*n=9216*).

To estimate doubling times, we used an automated set-up to count cells in colonies expanding on the designed variations of a solidified synthetic grape must, achieving generally high precision (mean CV = 10.0%) (Figure S2C). Designed selection environments, except for nitrogen starvation, increased the cell doubling time (mean increase: 4.15 h) compared to absence of a selection although with variation across lineages that reflects strong lineage-specific growth-differences in seven out of eight niches (Figure S4A – right panel). Commercial lineages enjoyed no cell doubling time advantage over pre-commercial in any niche, but many lineages in both categories were generally slow growers, implying that historical adaptation to the background grape must medium has been limited in these strains (Figure. S4A, S4B).

We evolved the 9216 wine yeast populations over 30 consecutive ALE batch cycles and stored evolved endpoint strains as frozen stocks (Figure S2). Extinctions of 2351 populations (∼25%) mostly affected slower growing lineages and likely reflected ratchet-like error accumulation and fewer viable cells being transferred to new batch cycles as the ALE progressed (Figure S6A). Surviving ALE populations varied dramatically in their endpoint doubling time adaptation, and differences between selection environments could explain 45% of this variance (Figure 2C-D). This reflects that adaptability in some environments is constrained across wine yeasts, while being generally high in others. Indeed, nitrogen starvation and selection for better aroma production often impaired rather than improved fitness (growth rates). This is consistent with the high rates of extinction in these designed niches, whereas the other six environments tended to promote fitness gains, albeit to varying degrees. The doubling time adaptability of wine yeasts were particularly impressive in high sugar, and to a lesser extent in fructose-rich grape musts, with 30% of the populations achieving >80% doubling time reductions. The capacity of wine yeasts to rapidly improve when facing environments richer in sugar than typical grape musts (200-250 g/L) is consistent with limited historical exposure to such yeast niches (e.g. honey), and thus, they will generally adapt fast to the climate induced increases in the sugar content of mature grapes^50^.

### Highly parallel ALE enables selection of better adapted variants

General differences between wine yeasts explained 23% of adaptation variance, illustrated by the strain E11 often adapting fast (mean doubling time reduction: 1.15h) while strain T7 consistently reached or spiraled towards extinction (Figures 2D, E, S6B). Wine yeast adaptability therefore has a generic component that persists across selection environments. Some lineages are inherently more suited to ALE, and the commercial and pre-commercial lineages are equivalent in this respect (Figure S6C). However, more often than not, adaptability involved significant genotype-by-environment components (Figure S6D), as illustrated by T14 matching the general adaptability of D10, but D10 adapting much better to fructose use and T14 much better to vitamin scarcity (Figure 2E). Predicting wine yeast adaptation in a particular ALE environment is therefore likely to be challenging, and this is further underscored by the best adaptation predictor, low initial fitness, accounting for no more than 36% of adaptation variance (Pearson, *r*=-0.61) (Figure 2F, left panel). An unbiased high throughput ALE platform based on the power of numbers circumvents this need for predictability by allowing a simple post-experiment selection of the most improved strains for further development. The value of such an approach is illustrated by the fact that the most adapted lineages reduced their doubling time 1.1-3.8x more than the average lineage in each environment (Figure 2C, inset) and often ended up among the fastest growing (Figure 2F, right panel). The latter underscores that ALE not only quenches defects in low-fitness lineages, transforming these into average wine yeasts, but imbues them with a fitness that is much superior to what is typical. This is illustrated by the gains of D10 in a fructose use environment and of E11 when selecting for relaxed nitrogen catabolite repression (NCR).

The divergence of populations evolving as replicates further amplified the value of the ALE parallelization, accounting for nearly 70% of adaptation variance within environments (Figure 2D). The most adapted ALE replicate of each parent (1.1-1.9x) consistently reduced its cell doubling time more than the average ALE replicate for that parent and for the best adapting parents (average of replicates) in each environment, this advantage of the best adapted replicate was even larger (1.1-6.6x greater doubling time reduction than the mean replicate) (Figure 2B and Figure S5). These benefits of ALE replication depended strongly on selection environment and genetic background, with selection for a relaxed NCR, tolerance to high sugar, and glutathione production, and lineages E12, G12 and M2 promoting a particularly large variation in adaptation between replicates (Figure S6D, bottom panel). Overall, we conclude that highly parallelized ALE improves the adaptation outcomes by expanding the numbers of ALE lineages and replicates, with both improving the chances of obtaining highly adapted variants.

### Parallelizing ALE empowers selecting against unwanted side effects

Pleiotropy, genetic linkage and neutral genetic drift means that ALE populations acquire traits other than those desired, often limiting their industrial usefulness. We measured side effects by cultivating the start and endpoints of 5760 evolved populations in 18 carbon or nitrogen limited environments and estimated the doubling time change that adaptation had caused. We found side effects on carbon and nitrogen metabolism to be common, with the mean population significantly (Student’s t-test, *p*=0.01) changing its cell doubling time in 15 out of 18 carbon or nitrogen limited environments. These side effects tended to be substantially weaker than adaptations, and they enhanced and impaired growth equally often and were decidedly non-random (Figure S7A). Thus, the capacity to grow under carbon restriction, particularly when provided as galactose, mannose, or maltose, often improved despite the absence of direct selection for use of these carbon sources (Figure 3A). In sharp contrast, the capacity to grow under nitrogen restriction, in particular when using aspartic acid, valine, and urea as nitrogen sources, consistently deteriorated across selection environments. This dichotomy likely reflects stronger indirect selection on a fast carbon metabolism in the carbon-rich background of synthetic grape must.

Next, we grouped populations based on evolved side effects using hierarchical and t-SNE clustering and found populations adapting to the same selection environment often acquiring similar side effects (Figure 3B, 3C). Thus, each selection environment allowed adapting wine yeasts to evolve a few (2-5) distinct sets of side effects, or syndromes, with each set being mostly private to one selection environment. Syndromes appeared to be universally available, as on average 96% of wine yeasts could acquire any given set of side effects (Figure S7B). However, different parent strains had very different propensities to acquire a particular syndrome in any given ALE niche. The strong influence of genetic background on the evolution of traits not under direct selection resulted in some wine yeasts tending to evolve more desirable sets of side effects than others, in the sense that these syndromes represented faster growth in a wide range of environments (Figure S7C). This tendency had a generic component, with e.g. M11 generally evolving fast growth side effects and G2 generally evolving slow growth side effects, regardless of ALE niche (Figure 3B, S7C). However, more often than not the founder strain evolving the best and worst sets of side effects varied across ALE niches. Given that predicting side effect evolution across genotypes is likely to be immensely challenging, agnostic implementation of ALE with many lineages in a highly parallelized platform therefore offers better chances of obtaining at least some ALE strains that are unburdened by unwanted side effects.

### *MEP2* and chromosomal mutations drive wine yeast adaptations to NCR relaxation

We sequenced the DNA of 26 fast adapting populations and called single nucleotide and structural variants with a substantial frequency in each population relative to its respective parent strain (Table S6). In the case of selection for NCR relaxation (arginine, proline consumption), four out of seven populations acquired amino acid changes in Mep2, implicating these mutations as drivers of this ALE adaptation (Figure 4A). Mep2, a high-affinity ammonium permease, also serves as main entry point for the ammonium analogue methylamine, which we used to activate the NCR intracellularly and to repress the use of arginine and proline, the only sources of usable nitrogen present in this ALE niche. Adaptation to this repression is therefore likely to have been driven by loss of Mep2 function and reduced methylamine uptake, NCR relaxation, and faster use of the arginine and proline. One of the point mutations, W275stop, is a non-sense mutation, consistent with this explanation (Figure 4B). An in-frame deletion of P465 in the auto-inhibitory domain of the cytoplasmic tail^51^ of Mep2 may result in constitutive Mep2 auto-inhibition and closure, with similar effects. We note that, while wine yeasts in ammonium-rich grape must initially take up ammonium through Mep1 and Mep3, Mep2 takes over when ammonium concentrations fall^32^. Loss of Mep2 function is therefore likely to result in slower ammonium uptake at later stages of wine fermentation, and consequently, to earlier NCR relaxation and more extensive uptake of the abundant proline and arginine, which is an industrially desired wine trait.

We found few other point mutations or small insertions/deletions in adapting populations, and these never affected the same protein (Table S7). Instead, all populations carried multiple aneuploidies, unbalanced chromosomal rearrangements and Loss-of-Heterozygosity (LOH) (Table S8). These were often extensively shared across populations, consistent with being selected for (Figure 4C). Some, e.g. the Chr XV duplication consistently emerging in a high sugar environment, were near private to specific ALE environments, suggesting niche-specific adaptations. Others, e.g. the Chr I and Chr XIV aneuploidies emerging in high alcohol, NCR relaxation, and glutathione production ALE niches, were shared across selection regimes and likely reflected adaptation to the synthetic grape must background medium - and potentially the good carbon us/poor nitrogen use common side-effect feature. Thus, although the underlying genes and mechanisms are challenging to pinpoint, many of our ALE adaptations were likely driven at least partially by large, recurring copy-number variations, consistent with earlier ALE observations^19, 20^.

### ALE populations express desired traits in larger liquid cultures

Because the physiology of yeast in larger liquid cultures differs from that of small colonies expanding on top of a solid matrix^52–55^, we probed whether ALE strains presented the wanted beneficial properties when shifted to larger liquid cultures (Figure S8A-B). We first tested whether production of the antioxidant glutathione, which protects cells against oxidative damage and can be secreted to also protect the grape must^56^, was present in adapted populations growing in 40 mL liquid cultures in closed vessels. Because we could only indirectly select for glutathione production, by supplying the glutathione precursors glutamate, cysteine, and glycine as the only nitrogen sources and challenging cells with the oxidant diamine, we expected many of the populations to have adapted by other means. Nevertheless, we found three of eighteen tested populations, M4 (15,28), E5 (15,39) and E6 (31,30), to have increased their glutathione production by between 35% to 62% when cultivated in liquid (Figure S9A), with the best performer M4 (15,28) producing ∼13 nmol glutathione/mg dry weight of biomass.

We next explored whether ALE populations selected to relax NCR, and the use of arginine and proline even in the presence of a preferred nitrogen source, express this property when shifted to 80 mL liquid cultures. We selected 11 ALE populations having evolved faster growth on the NCR selection medium and cultivated them in liquid, mostly anaerobic cultures, using ammonium as NCR inducer. We found E11 (21,4) and E3 (17, 32) to consistently take up less (46 and 61% respectively) ammonium than G4, a top pre-commercial strain, while consuming at least as much as sugar (Figure S9B). Endpoints of both these ALE populations also finished fermentations faster than their start points, when growing on arginine and proline as sole nitrogen sources and using methylamine as NCR inducer. This suggests NCR relaxation and concomitant faster use of arginine and/or proline (Figure 5A). We therefore tracked the assimilation of these amino acids in these ALE cultures by HPLC, finding that while proline uptake remained unchanged, net arginine uptake had improved. ALE populations achieved this better arginine use by avoiding the early loss of arginine to the environment suffered by their founders (Figure 5B). Yeast store arginine in the vacuole as a nitrogen reserve^57^, and if a preferred nitrogen source is encountered before this stored arginine has been mobilized, it is instead actively exported^58^. Our results are therefore consistent with some ALE populations reaching the desired improved arginine consumption by relaxing NCR and avoiding early arginine export to the grape must.

**Figure 5.**
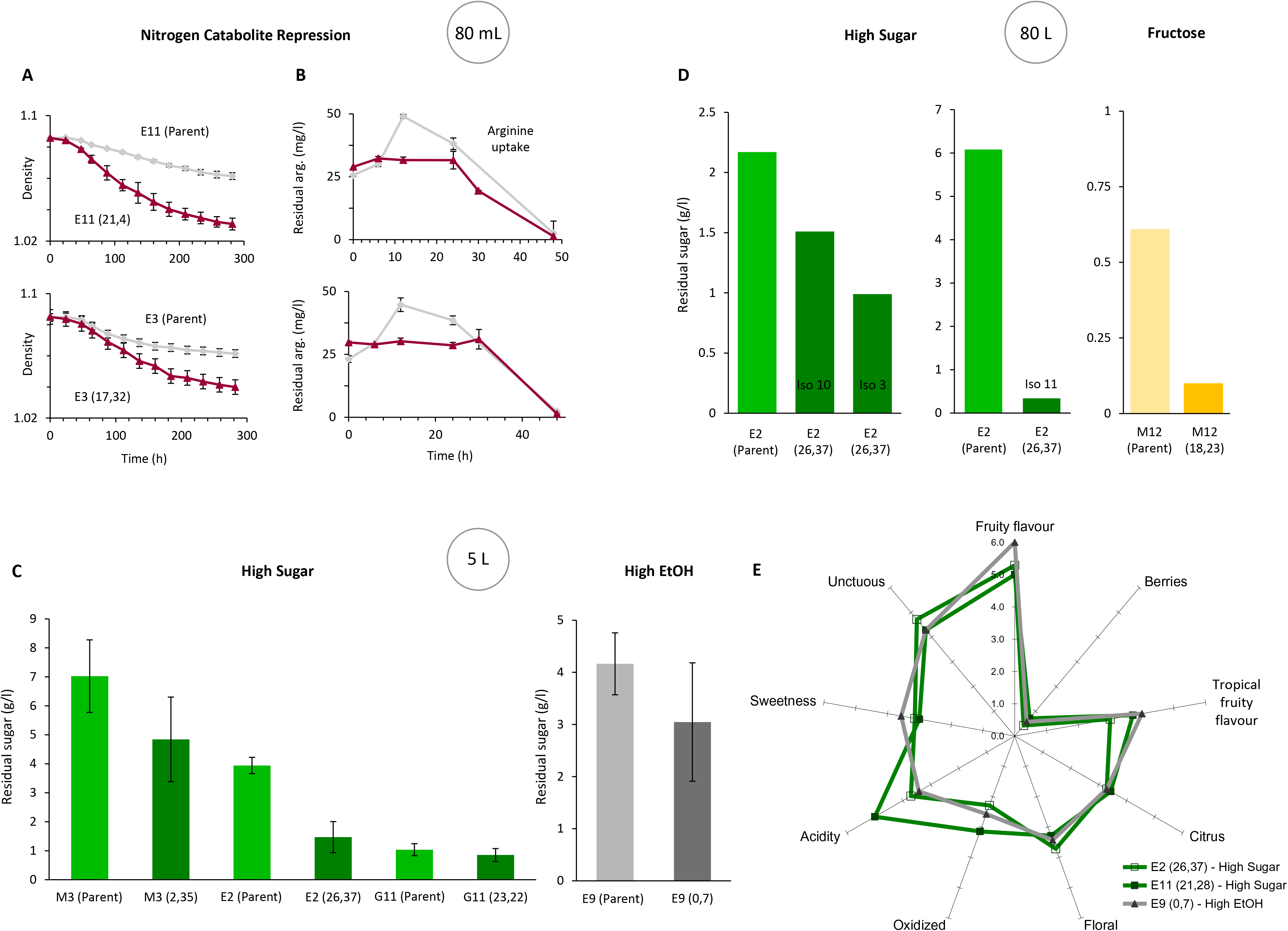
ALE populations retain adaptations in larger cultures. **(A-B)** We cultivated two ALE populations selected for NCR relaxation and with consistently low ammonium uptake (see arrows in Figure S9B) on selection medium in larger, liquid cultures. Founder lineages are shown as references. Numbers in parenthesis: adapted colony ID. **(A)** We tracked the fermentation (carbon dioxide production) by measuring the density of cultures. Error bars: SEM (*n*=3). **(B)** We tracked the net arginine uptake in cultures by measuring the residual arginine in the selection medium by high-performance liquid chromatography (HPLC). Error bars: SEM (*n*=3). **(C)** We cultivated clones drawn from ALE populations evolved for high ethanol tolerance and three evolved for high sugar tolerance in 5 L actual grape must and compared the fermentation capacity (residual sugar after 32 days) to that of their founders. Error bars: SEM (*n=3* clones). **(D)** We cultivated one clone drawn from ALE population M12 (18, 23) evolved for fructose use and three clones from ALE population E2 (26, 37) evolved for high sugar tolerance in 80 L actual grape must and compared the fermentation capacity (residual sugar after 32 days) to that of their founder lineages. **(E)** Organoleptic properties of the wine produced by two ALE populations selected for high sugar tolerance and one selected for high ethanol tolerance was scored from 0 to 6 by a blinded taste panel of three experts. Numbers in parenthesis: adapted colony ID.

### ALE strains perform well in industrial grape must fermentations

We next probed whether evolved populations perform well in cultivation scales that approach wine production conditions and in actual, rather than synthetic, grape must. For this, we selected eight ALE populations with good growth in high alcohol, high sugar, or high fructose ALE environments. We first tested whether populations adapted to high alcohol in the form of 1.3% n-butanol, which resembles ethanol but is less volatile and remains in the solid medium^2425^, also grew better when exposed to high ethanol in 40 mL liquid cultures. We found three of the six tested ALE populations to grow both faster (5-37%) and to higher (78-137%) cell yield in 8% ethanol than their founder strains (Figure S9C). Two of these, T3 (13,35) and T3 (15,35), also fermented the sugar ∼14% more efficiently than their founders in absence of added ethanol, and we selected both of these for wine cellar experiments (Figure S9D).

Next, we probed whether 18 ALE populations adapted to a high sugar content also fermented a high-sugar synthetic grape must better when cultivated as 40 mL liquid cultures. Five populations either left less sugar in the must at the end of the fermentation than their founders or produced more ethanol per sugar molecule consumed. We retained these five populations for wine cellar experiments (Figure S9E). Finally, we cultivated ten ALE populations adapted to fructose use, in liquid 40 mL fructose cultures also containing the glucose analog 2-deoxyglucose. We found one to grow much faster, reach a higher cell yield, and assimilate 35 g more fructose than its founder, and we retained this for the wine cellar experiments (Figure S9F).

We isolated single clones from each of the eight selected ALE populations, expanded these clonally to create large starters and inoculated the latter in 27% sugar White Grenache (GR) grape must recently harvested in DO Terra Alta (Spain). We cultivated four of the eight ALE strains, in triplicate, in 5 L cultures and 4, non-replicated, in 80 L cultures and tracked their capacity to ferment the grape must. Fermentation progressed without detectable defects in all eight ALE lineages, with six that were adapted to either high sugar, fructose use, or high ethanol, consuming up to 2% more of the sugar than their founders (Figure 5C, D). We evaluated the visual and organoleptic properties of the wine produced by three of these ALE lineages using a blind taste panel of three experts and found them to give similar fruity, unctuous flavors (Figures 5E). Our highly parallelized ALE platform was therefore capable of evolving strains with a better performance that persists in industrially relevant fermentations and without defects in wine taste and smell.

## DISCUSSION

### Parallelization improves ALE outcomes by broadening the evolutionary search spectrum

We introduced a highly parallelized ALE platform for improvement of industrial microbes that is based on expanding ALE populations over many generations as colonies on top of a designed solid selection medium and accurately counting their cells. The platform allowed us to broaden the evolutionary search spectrum both by repeating ALE many times from a fixed genetic start point and to expand the number of different start points. Both improved the chances of obtaining ALE populations with much better growth in the ALE environment and reduced the risks of ALE burdening strains with poor growth side effects. This agrees with population-genetic predictions. New mutations emerge stochastically, and when rare, chance heavily affects their fates. Thus, some divergence between replicated ALE populations is expected. Genetic start-point variation amplifies this divergence, as the phenotypes encoded by new mutations often depend on other variants that are already present^59, 60^, such that the latter guides evolution down different evolutionary paths^61, 62^.

Arguably, the benefits of ALE parallelization depend on the specifics of the selection regime. In the current design, the relatively small population size (<3×10^7^ cells) means that strongly beneficial mutations will occur only rarely^63^. In larger ALE populations, the most advantageous mutations will occur more often and this should speed up adaptation, reduce variation in adaptation across replicated ALE populations, and consequently also diminish the benefits of ALE parallelization. However, the strongest mutations are often associated with side effects that impair growth in non-selection niches^64^. Moreover, they tend to be incompatible with each other^65^, leaving large populations stranded on suboptimal local fitness peaks and incapable of sustaining adaptation. Our current design, ALE of many small populations, may therefore have advantages that ALE of a single, or even many, larger ALE population will struggle to match. The empirical data from our study indicates an average rate of successful improvement (>2 fold improvement in growth rate) to be around 4*10^-5^ per generation (∼60 improvements among 6664 lineages, with circa 200 generations per lineage). This is remarkably low - a low throughput ALE with tens of parallel lines of a single strain would in comparable environments require multiple years for obtaining a single successful clone - attesting to the advantage presented by massively parallel experimental evolution demonstrated here.

Wine yeasts often inherit all, or the lion’s share, of their genome from the Wine/European *S. cerevisiae* clade^48, 49^ from which our parent strains all descend. Despite their close relatedness (mean pairwise nucleotide diversity, π: 0.00068), they nevertheless differed markedly in adaptation and side effects, in line with differences in even a single gene often altering evolution^66^. The power of highly parallelized ALE will arguably increase with greater diversity among genetic start points and inclusion of rare wine yeast with admixed or hybrid genomes or carrying DNA introgressed or horizontally transferred from other clades or microbes^67, 68^, may in this context be particularly valuable. We note that an alternative ALE design pooling strains to generate diverse starting populations^69^ will not exploit the initial genetic diversity to the same extent because strong early selection on the best pre-existing variants will restrict the subsequent *de novo* mutation-based evolutionary search^70^. And meiotic recombination offers no easy workaround^71, 72^ as domestication has impaired the sexual life cycle of most industrial budding yeast strains^73^.

Despite the large body of literature reporting a decelerating adaptation as populations become fitter^29, 74–76^ because of the weaker effect of new mutations in fitter backgrounds^77–80^, diminishing returns only explained a minor fraction of the variation in doubling-time gains among our ALE populations. Thus, our platform improved ALE outcomesacross the parental strain performance spectrum, and the ALE populations with the largest doubling-time gains often ended up being among the best performers, and superior to all founders. The moderate influence of diminishing-return adaptation is reassuring, as it demonstrates that ALE is not restricted to compensating for defects in poor-performing lineages, but can also improve the best performing wine strainss, and thus, has potentially broad versatility.

Apart from the consistent point-mutation based inactivation of the ammonium and methylamine importer Mep1 during selection for NCR relaxation, aneuploidies emerged as the most likely drivers of the majority of ALE adaptations, consistent with what has been observed in *S. cerevisiae* lab strains^81–88^ and in drug-treated fungal pathogens^89, 90^. Because of their recurrence across populations and reported high degree of pleiotropy^91–93^, aneuploidy also appears to be the main etiological agent behind the ALE emergence of distinct syndromes, and in a particular of the meta-syndrome of “good carbon, poor nitrogen use”. It is attractive to speculate that the latter derives from a rerouting of carbon fluxes from amino acid anabolism, which originate in pyruvate and TCA cycle intermediates, to fermentation, and that this re-routing is favorable under amino acid excess when carbon-use limits growth, but detrimental under nitrogen deprivation. The ancient duplication of the *S. cerevisiae* genome is held to explain its good use of fermentable carbon^94^, supporting that concerted amplifications of glycolytic genes is capable of driving such evolutionary shifts. As our synthetic grape musts contain >10x the fermentable carbon of standard laboratory nutrient medium, selection for good use of this carbon should be strong in most of our designed ALE environments. Implicit in this conjecture is that wine yeasts are far from optimally adapted to carbon rich environments, being constrained by the strong selection for growth under nitrogen deprivation^95^.

Our ALE platform generated improved wine strains that often expressed the industrially desired traits in larger, liquid cultures and produced good quality wines. Hence, not only does it bring evolutionary engineering into the realm of high-throughput science but it also opens a new fast-track lane for optimizing microbes for industry-desired traits. And because the platform works well with microbes covering broad swaths of the tree of life, including bacteria^96^, we expect it to be of substantial value across many industrial sectors.

## Supplementary Figure Legends

**Figure S1.**
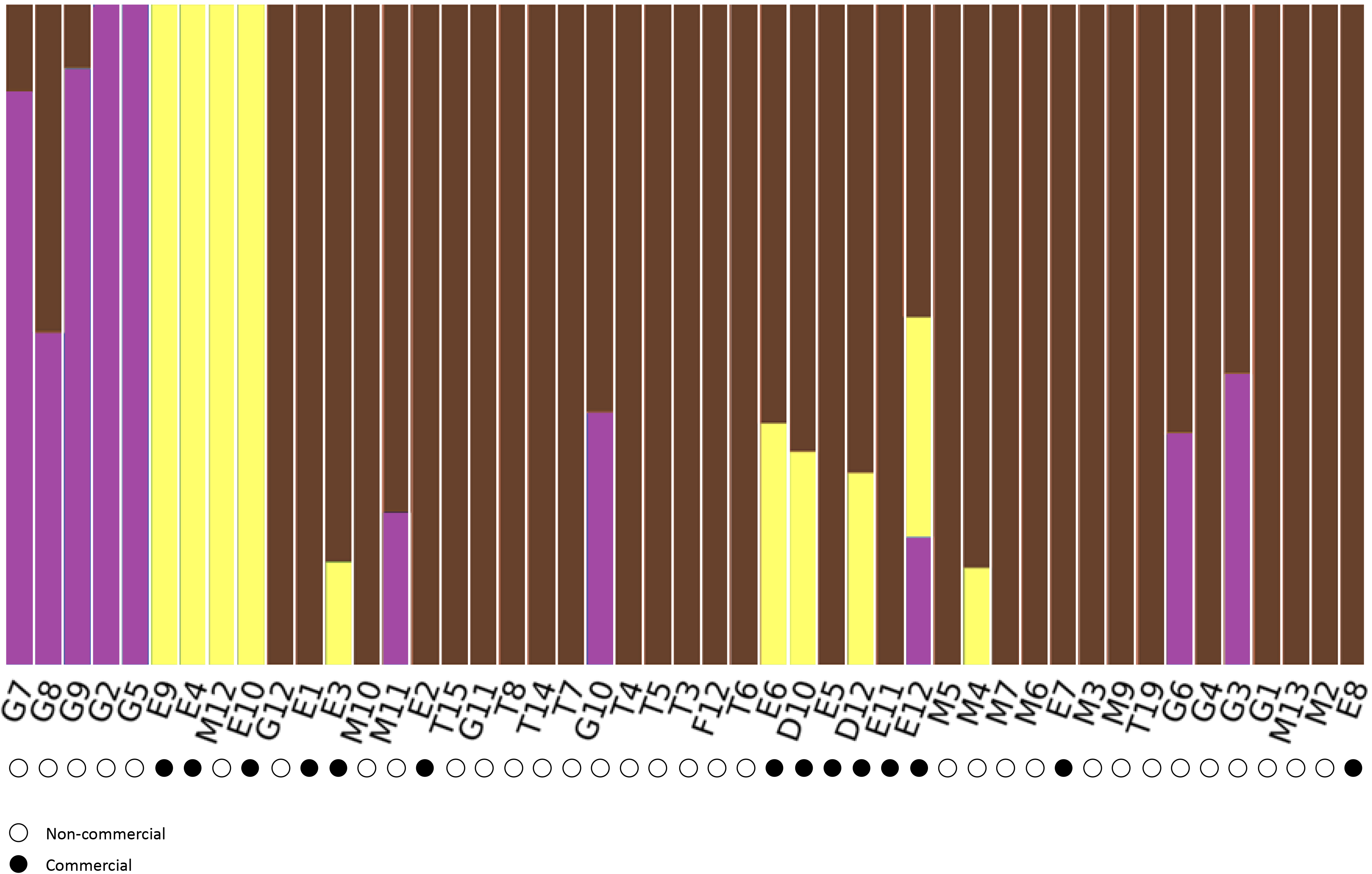
Population structure of parent wine strains, before ALE. We sequenced the genome of 48 commercial (●) and pre-commercial (○) euploid diploids wine yeast lineages (Table S1) using Illumina and identified their population structure^49^ based on 42,599 SNPs called against the EC1118 wine strain reference genome. We varied the number of populations (*K*) between 1 and 10 and show the structure for *K*=4 (colour).

**Figure S2.**
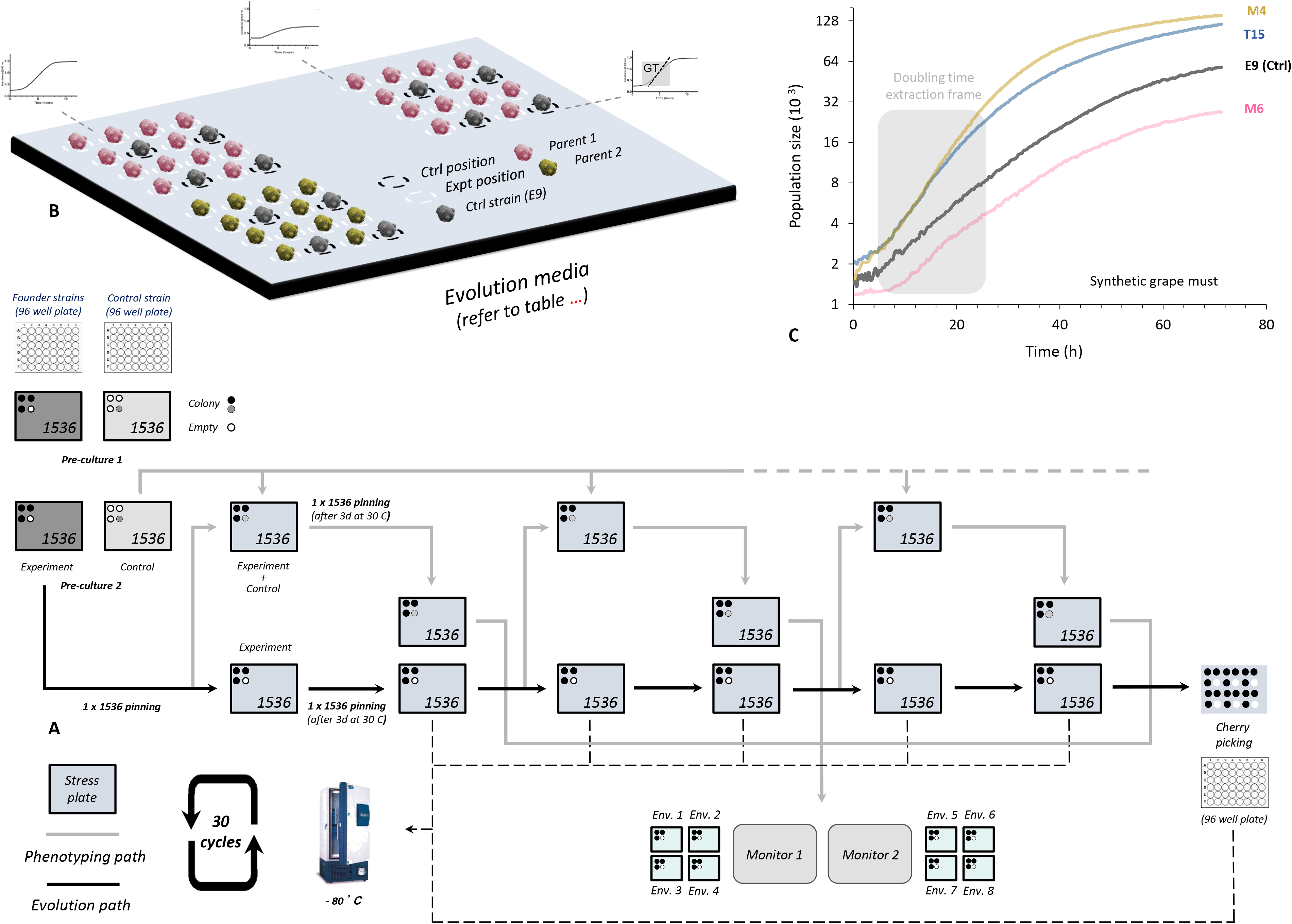
Parallelizing wine yeast ALE. **(A)** 48 commercial and pre-commercial wine yeast strain (Table S1) were ALE evolved as replicated (*n=24*) asexual populations over 30 growth cycles. ALE populations were cultivated (black arrows = evolution track) as colonies on eight synthetic grape media designed to select for traits desired by the wine industry (Table S2 and S3). We generated 48 founder populations from single clones of each founder strain (not shown), and replicated these 24x to generate a 1152 colony array cultivated on synthetic grape must. Colonies were stored at −80°C in 96 arrays in 20% glycerol. Frozen stocks were revived on synthetic grape must and transferred to evolution plates representing the eight selection regimes. The 9216 ALE populations were passed through 30 cycles of growth, sampling and transfer (*n=∼10^5^* cells) to fresh plates. We stored the cycle 30 end-point of each population as a frozen ALE record before reviving and cultivating these in 1536 format, while counting cells in each growing colony (grey arrows=phenotyping track). The 1152 cycle 0 start points were revived and cultivated in parallel, on separate plates. Populations were cultivated as multiple replicates (*n*=2-4; on separate plates) in each of the eight designed selection environments and in the 18 nitrogen or carbon limited side effect environments. A fixed control was introduced and cultivated in every 4^th^ position on every plate and used to control for systematic growth variation between and within plates. We extracted cell-doubling times from high quality growth curves, log_2_ transformed and normalized these measures to those of the 384 fixed controls on the same plate. The output ∼0.5 million growth measures is reported in Data S1. **(B)** A zoom-in view of one experimental plate, showing the arrangement of experimental and fixed control populations. **(C)** Growth of wine yeasts (color) growing on synthetic grape must. Black = fixed control (parental strain E9). Grey field: time window in which the cell doubling time was extracted

**Figure S3.**
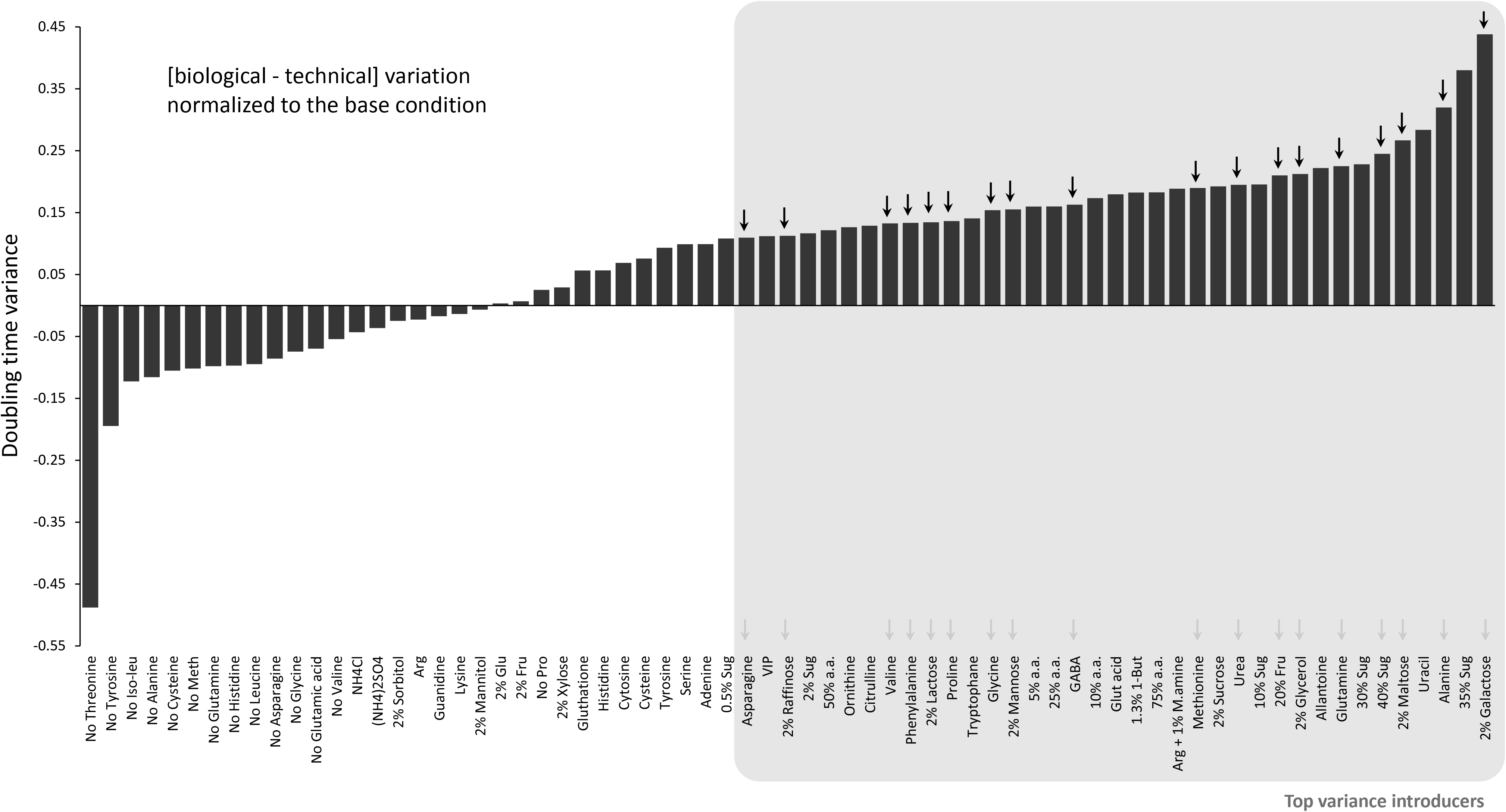
Growth variance of parental wine yeasts in synthetic grape must. The 48 parental wine yeast strains were cultivated in 70 synthetic grape must, distinct in C and N sources, and their log_2_ doubling time relative to a fixed control (strain E9) was estimated (*n*=24 replicates). Technical variance (across replicates) was subtracted from biological variance (across 48 strains) and then normalized to the variance in standard synthetic grape must. Arrows: selected environments for side-effect experiment (Figure 3).

**Figure S4.**
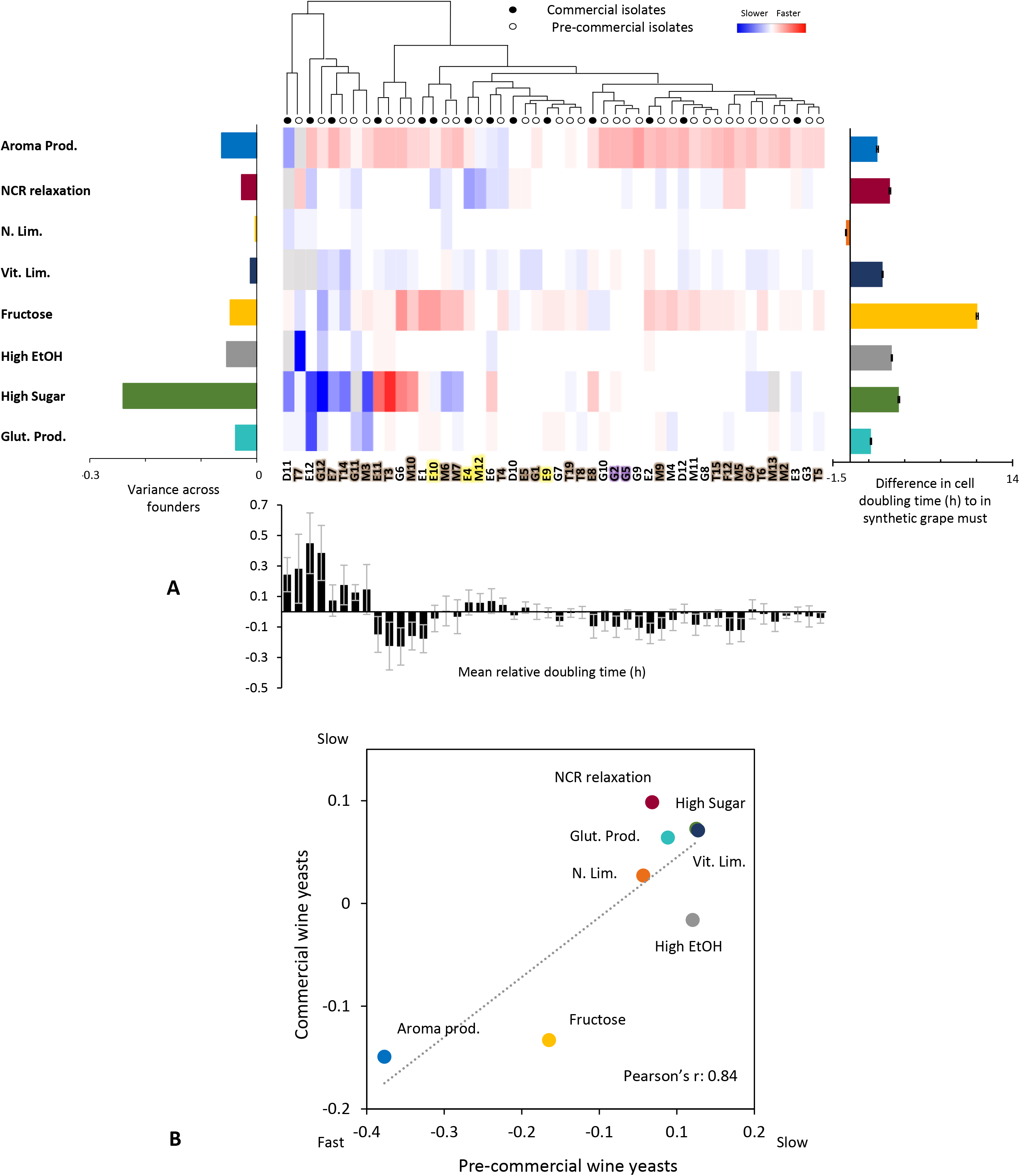
Strength of ALE selection. Cell doubling times of wine yeasts in selection environments (Table S2 and S3). **(A)** *Central panel (heat map):* Cell doubling times (mean: *n*=24) of each wine yeast normalized to the fixed control (color). No color: fixed control. Wine yeast names are colored based on population (see Figure S1). *Upper panel:* Hierarchical clustering of wine yeast based on similarity in cell doubling times, using Pearson’s *r* and strain averages (for groups). *Left panel*: Variance across wine yeasts (*n*=48) in mean cell doubling time (*h*). *Right panel*: Strength of selection, estimated as mean (*n*=48 wine yeast) difference (*h*) in cell doubling time in an environment as compared to in synthetic grape must (SGM). Error bars = SEM. *Bottom panel*: Some wine yeasts are general slow growers, reflecting limited adaptation to synthetic grape must. Mean of normalized cell doubling times for each wine yeast, across all selection environments. Error bars = SEM (*n=8* environments). **(A)** Commercial (*n=15*) and pre-commercial (*n=33*) wine yeast grow equally well in all selection environments. Mean cell doubling times normalized to the fixed control are shown. Broken line = linear regression.

**Figure S5.**
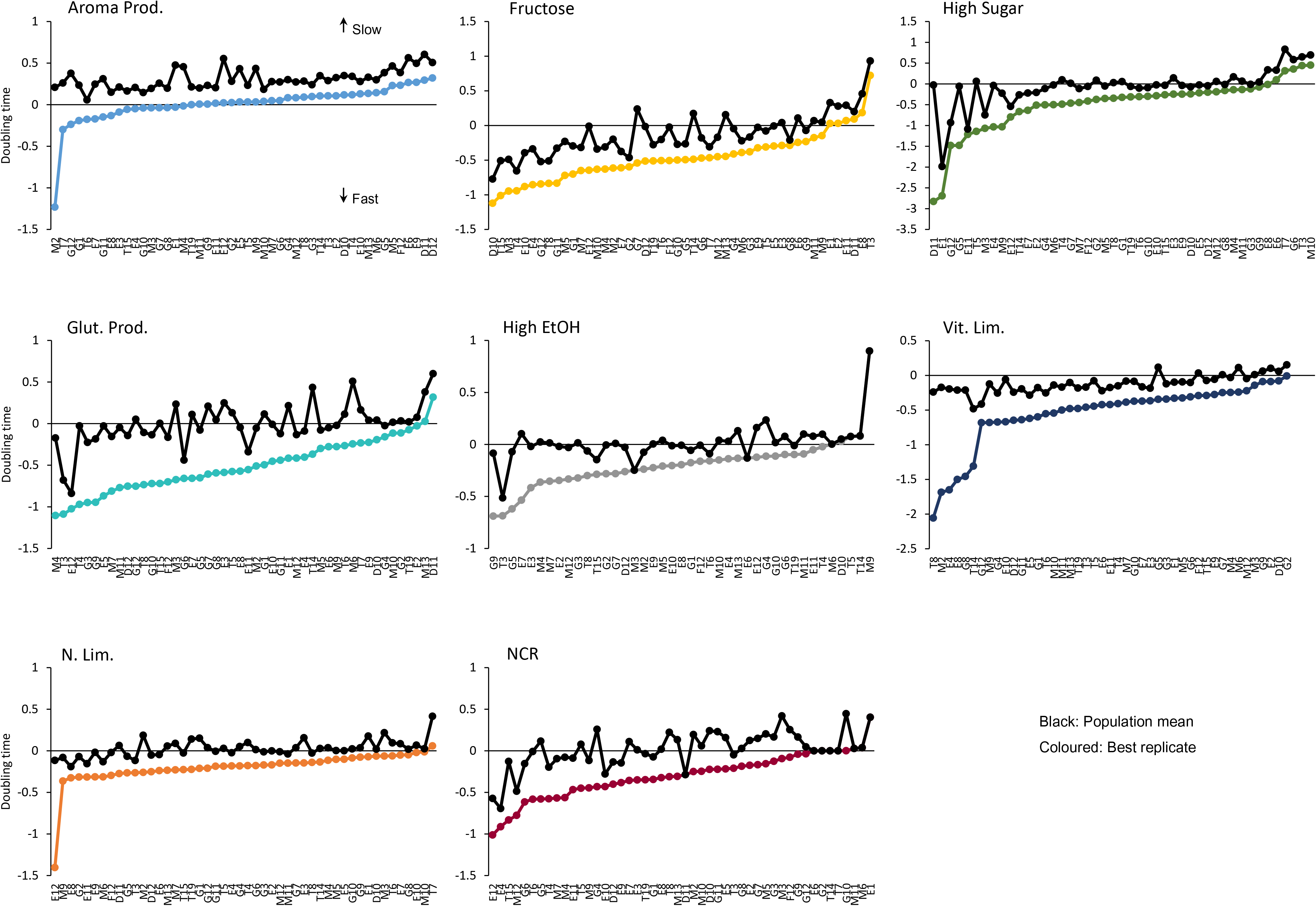
Replication resolution of adapted wine yeasts. Doubling time line chart of the best adapted replicate (coloured) and population mean (average of all replicates: black) of each lineage in each environment. Error bars = SEM (*n=2-4* for the best replicate and *n=24* for the population)

**Figure S6.**
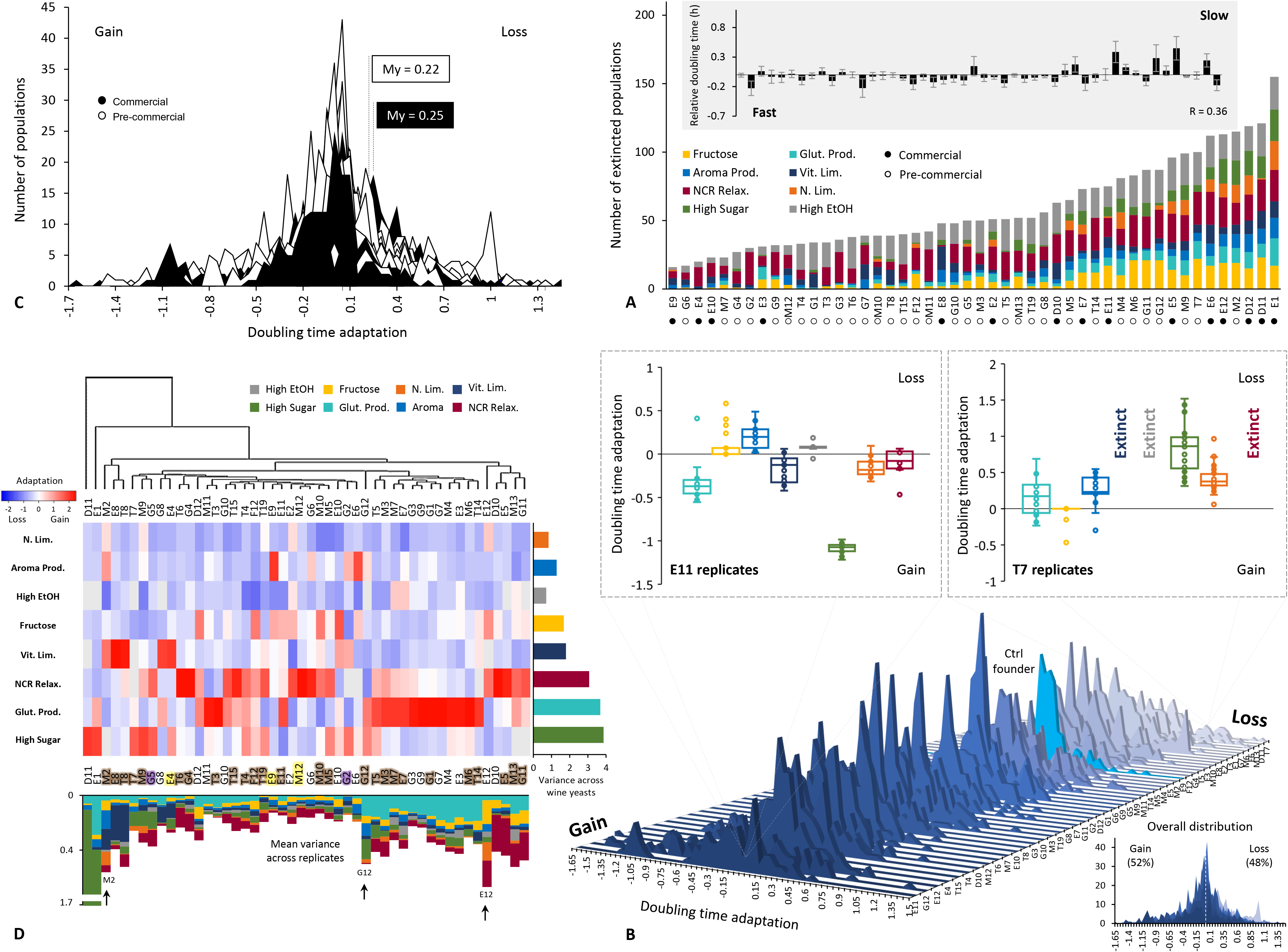
Adaptation of ALE wine yeasts. **(A)** Extinct populations: stacked bar plot of number of extinct populations, in each environment. Inset: mean of (log_2_, normalized) cell doubling time. Error bars: SEM. **(B)** Adaptation (*n*=24, extinct populations excluded) of wine yeasts E11 (left panel) and T7 (right panel) in selection environments (colour). Box: interquartile range, black line: median: whiskers 1.5x interquartile range, outliers: populations outside interquartile range. **(C)** Histogram of adaptation for all commercial (black, *n=2880)* and pre-commercial (white, *n=6336*) wine yeast populations, in all selection regimes. Extinct populations are excluded. **(D)** Adaptability has a genotype-by-environment component. *Central panel (heat map):* Adaptation (mean: *n*=24) of each wine yeast. No color: no adaptation. Wine yeast names are colored based on population (see Figure S1). *Upper panel:* Hierarchical clustering of wine yeast (Pearson’s *r*, averages used to cluster groups) based on similarity in adaptation across eight selection regimes, using *Left panel*: Variance in mean adaptation between wine yeasts, in each environment. *Bottom panel*: Variance in adaptation between replicated populations (*n*=24), for each wine yeast in each environment. Lineages mentioned in text are indicated (arrows).

**Figure S7.**
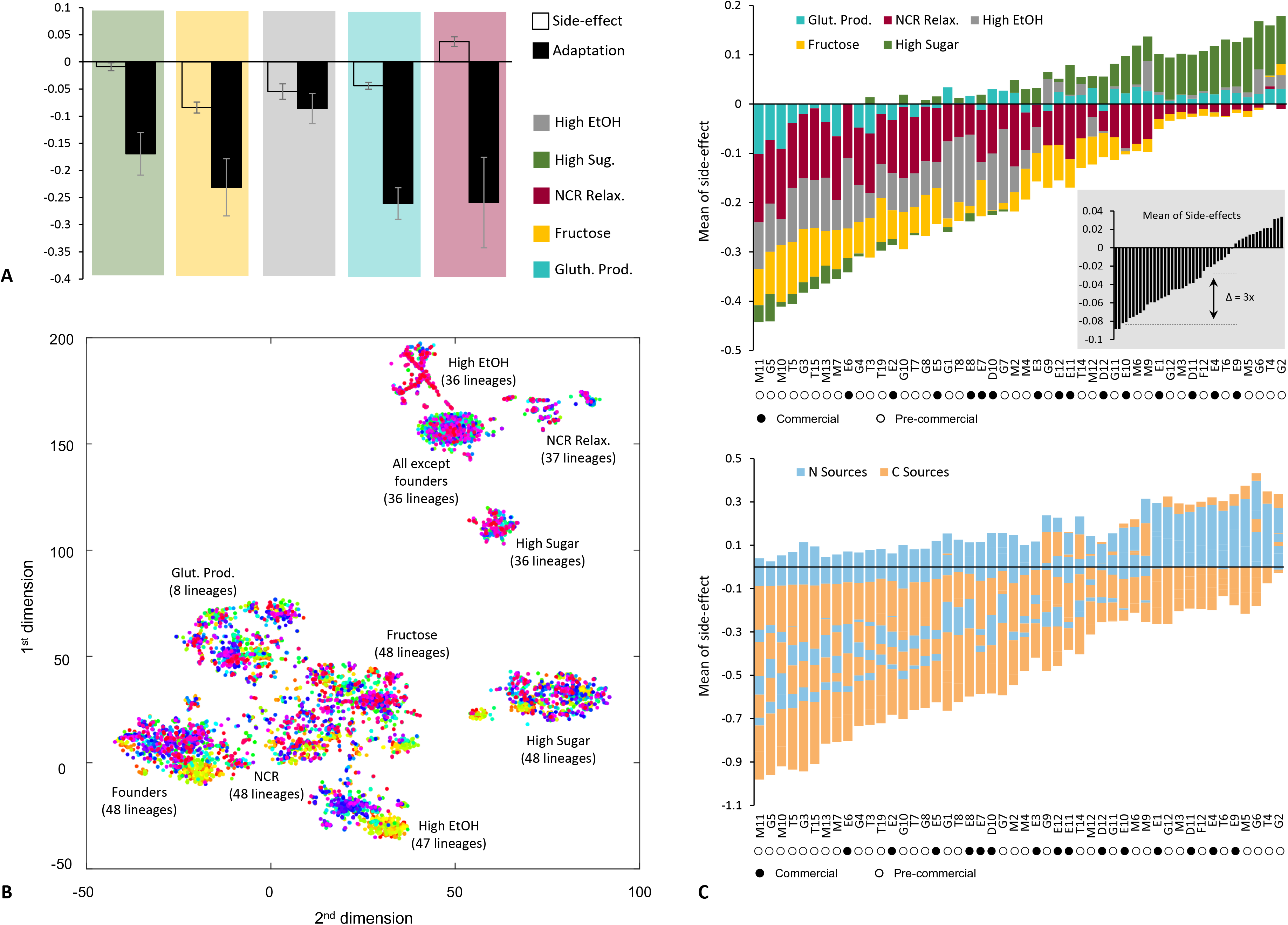
Side effects evolved by ALE wine yeasts. **(A)** Side effects (top bar) of evolution are smaller than adaptations (bottom bar), in all environments. Means across all side effects (*n*=18), and across all populations (*n*=24) of all lineages (*n*=48) in each environment are shown. Error bars = SEM across lineages (*n*=48). **(B)** t-Distributed Stochastic Neighbor Embedding (t-SNE) clustering reducing the variance in side effects to two dimensions (*x*, *y*-axes). Each dot represents one population of one lineage in one selection environment, or one starting population. The clustering is identical to in Figure 3, but colour indicates lineage. For large clusters, the selection regime and the number of lineages in the cluster are indicated. Note: side effects do not cluster by lineage and each cluster contains representatives of almost all lineages. **(C)** *Upper panel*: some wine yeast lineages (left) evolve more desirable (faster growth) side effects than others (right). Stacked bar plot of means across all side effect environments (*n=18*) and all populations (*n*=24) for each lineage is shown. *Inset*: barplot of means. Colour = selection environment. Negative numbers: cell doubling time reductions. *Bottom panel:* stacked bar plot of means across all population (*n=24*) in each carbon or nitrogen environment.

**Figure S8.**
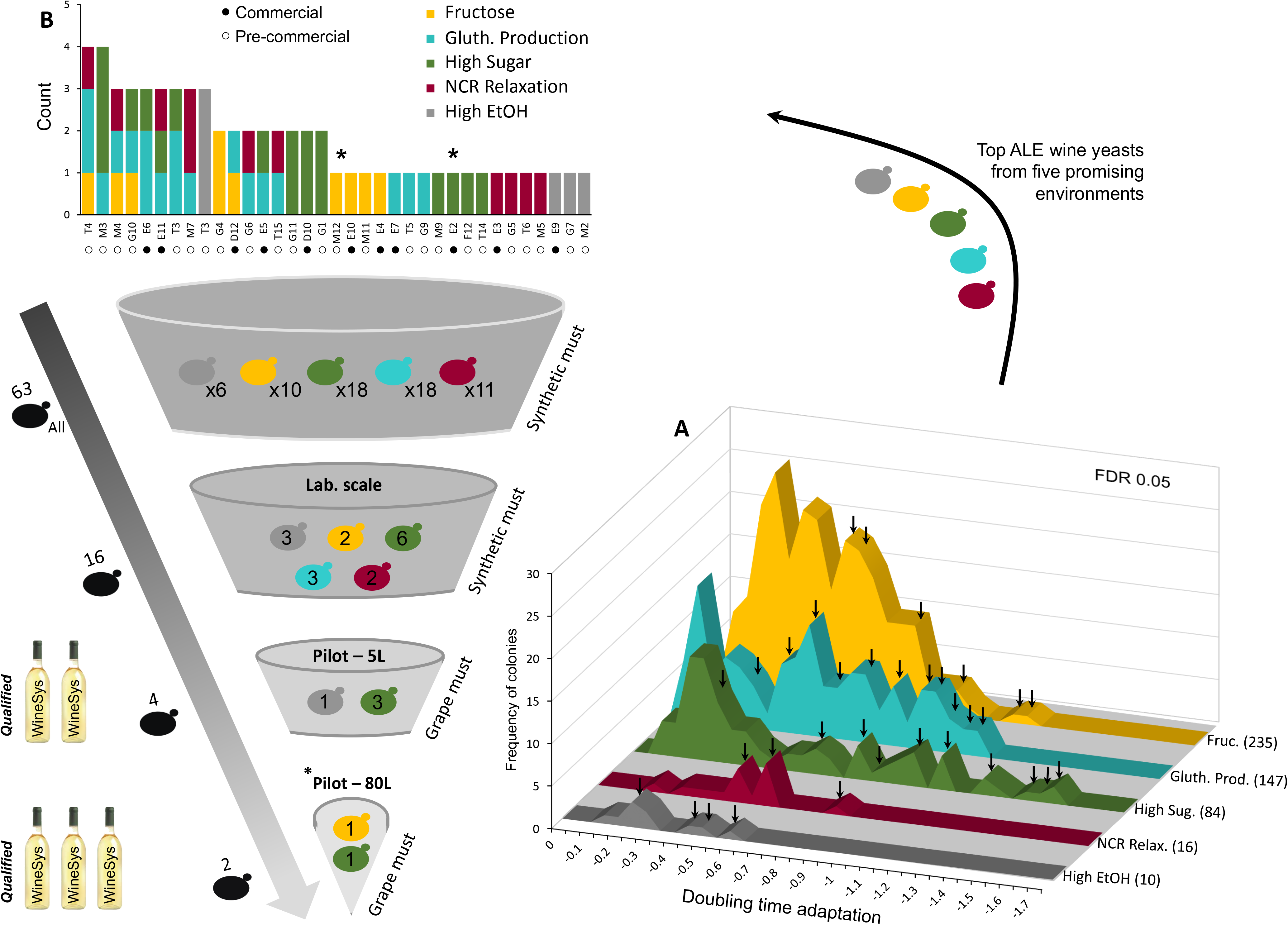
Selecting ALE populations for larger culture validation. **(A)** 3D histogram of adaptation of populations with shorter doubling time compared to their founders in selection regime (colour). Environments with little or no adaptation are excluded. Arrows: ALE populations selected for validation. In parenthesis: number of ALE populations with significantly different doubling times compared to the founder (FDR: q=0.05). **(B)** Bar plot of selected ALE populations. Asterisk: also cultivated in 80 L grape must +-cultures.

**Figure S9.**
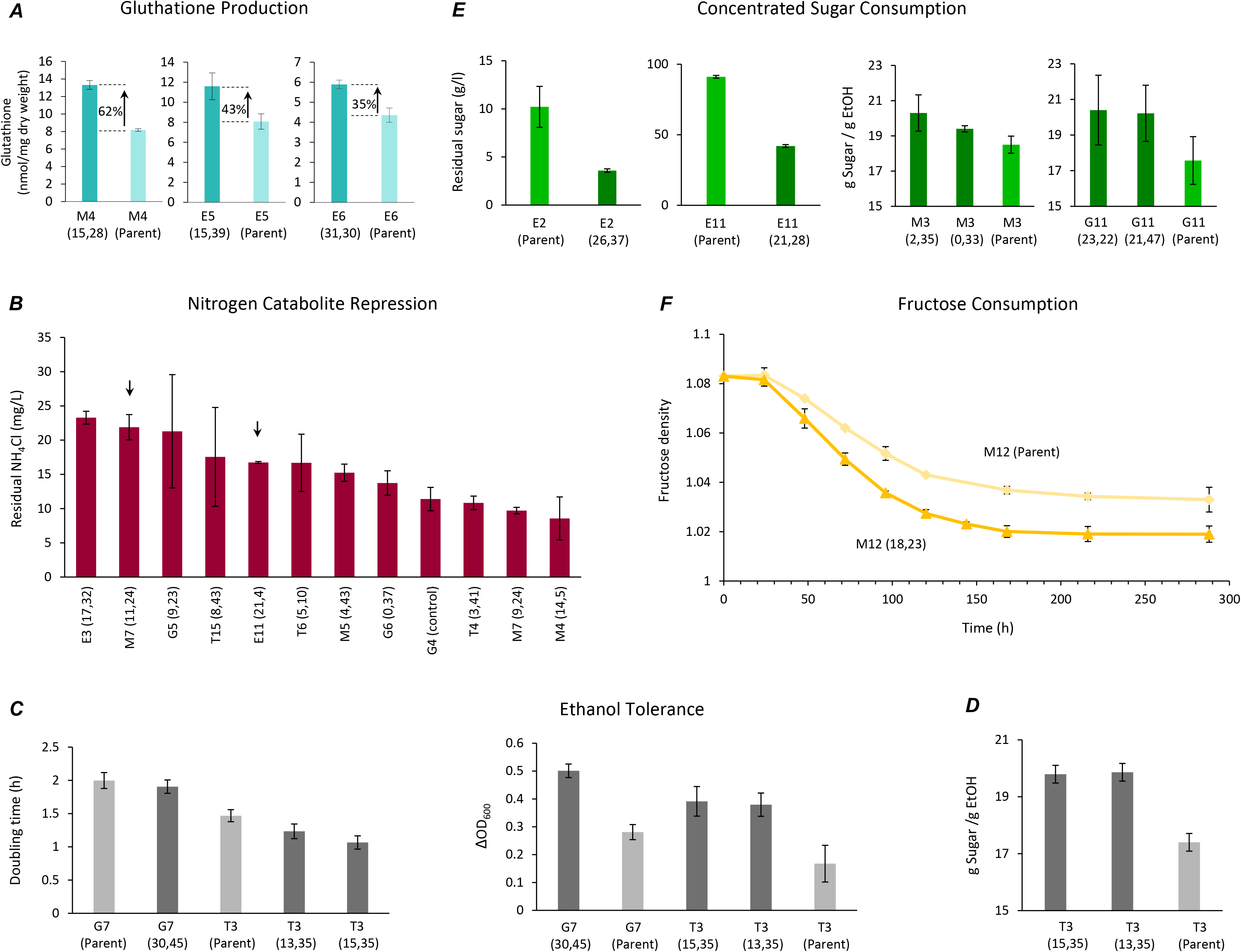
ALE wine yeasts retain adaptations in larger liquid cultures. **(A)** Three ALE populations selected for higher glutathione production produce more glutathione than their founder lineages in larger, liquid cultures. Total glutathione (nmol/mg dry weight biomass) at the end of a growth batch cycle (168h) is shown. Error bars: SEM (*n*=2). **(B)** 11 ALE populations with improved growth in the NCR relaxation selection regime was cultivated in liquid synthetic grape must cultures (250 mL) with ammonium (43 mg N at start) as preferred nitrogen source and the residual ammonium was measured at the end of fermentation (360 h). The founder strain G4, as the best performing pre-commercial wine strain, is shown as reference. Error bars: SEM (*n*=3). Arrows indicate ALE populations with consistently low ammonium assimilation. Note that non-ammonium nitrogen (total: 100 mg N at start) were mostly present as arginine and proline. **(C)** Six ALE populations, evolved for higher ethanol tolerance, and their founder strains, were cultivated in 40 ml liquid cultures in the presence of 8% ethanol. We tracked their growth continuously and extracted cell doubling times and cell yields. **(D)** We cultivated two ALE populations evolved for higher ethanol tolerance and expressing this trait in larger liquid cultures (see Figure S9C) in 50 mL synthetic grape must (*n=2-3*) and compared their fermentation efficiency, measured as the gram sugar consumed per ethanol produced, to that of their founder strain. Error bars: SEM (*n*=3) **(E)** 18 ALE populations, evolved for better growth in high sugar concentrations, were cultivated in 40 mL liquid high sugar grape must. We tracked the sugar consumption and ethanol production in each. Two ALE populations showed faster sugar uptake (*top panel*) than their founder strains and four showed more efficient fermentation (*bottom panel*; gram ethanol produced per gram consumed). Error bars: SEM (*n*=3) **(F)** One ALE population, evolved for better fructose use, was cultivated in a 40mL liquid fructose containing synthetic grape must in the presence of the glucose analog 2-deoxyglucose. We compared its sugar fructose consumption to that of the founder strains. Error bars: SEM (*n*=3).

**Table S1.**
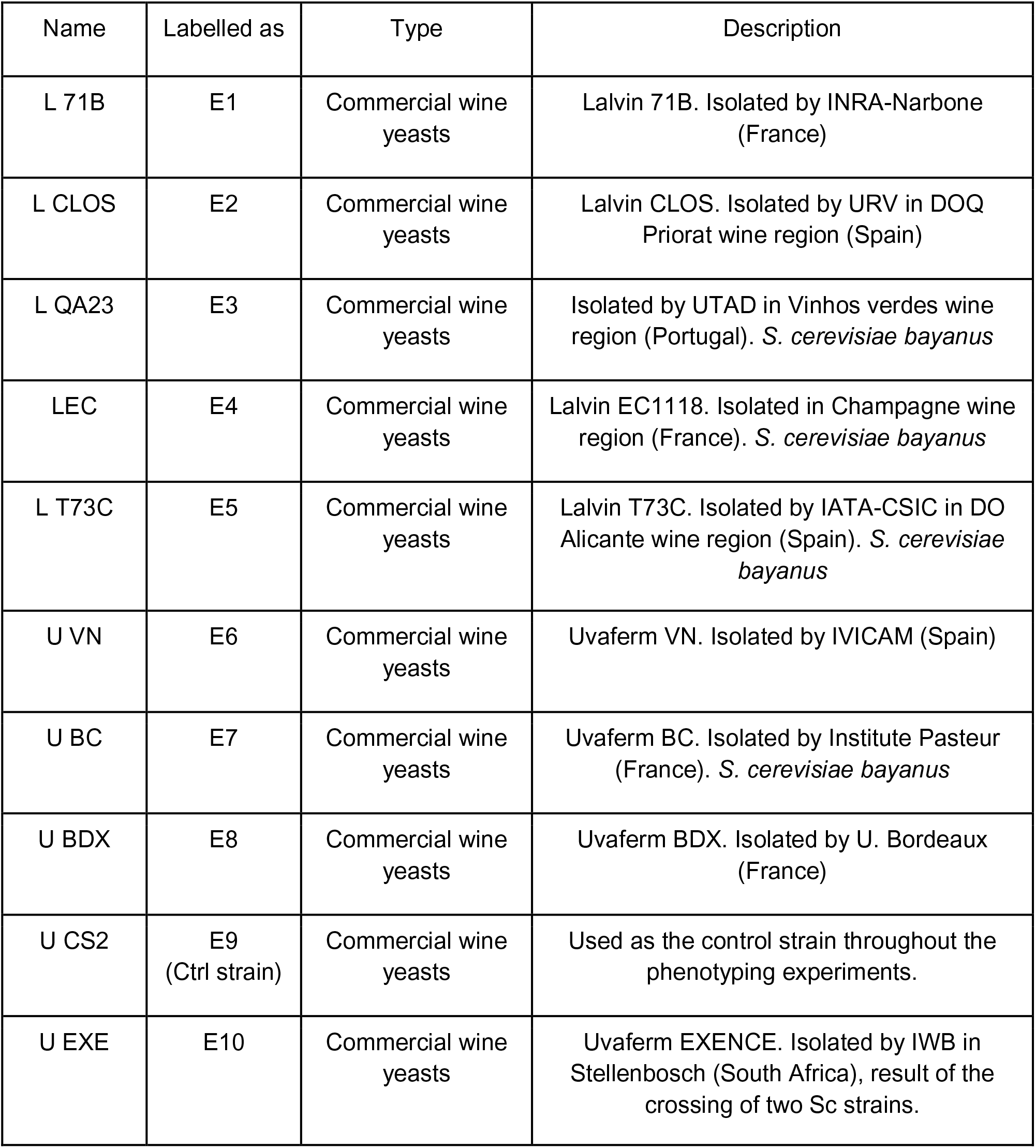
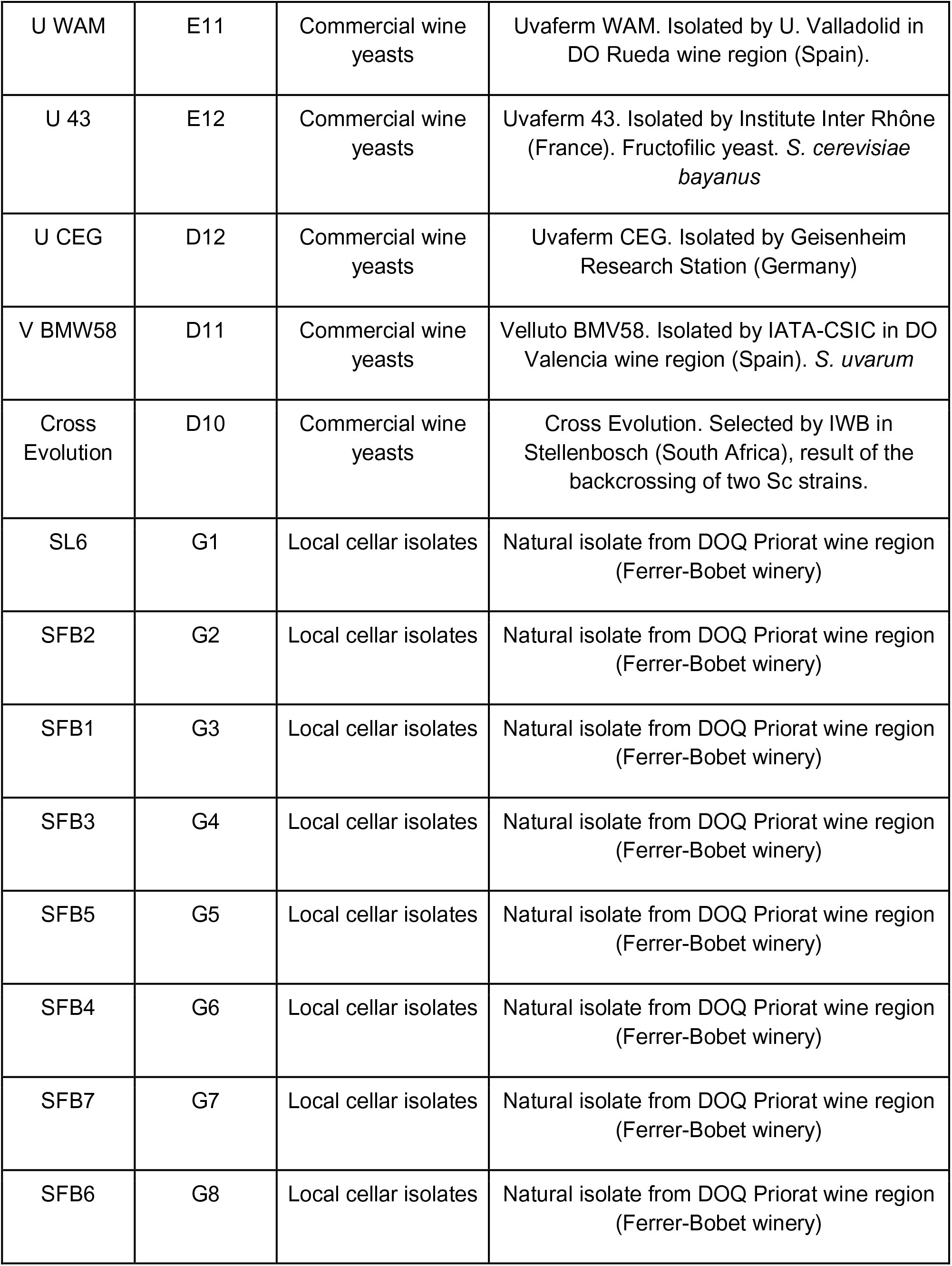
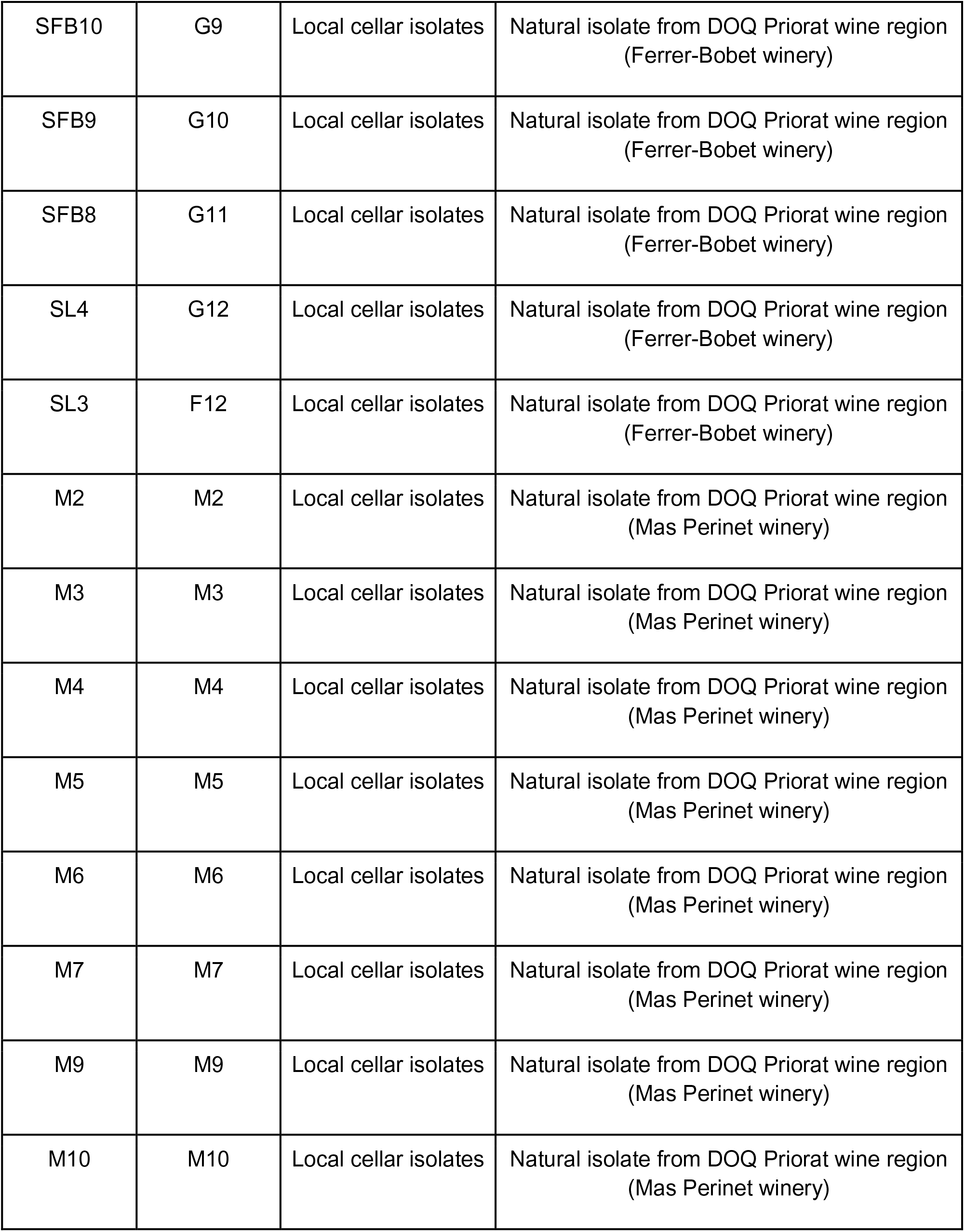
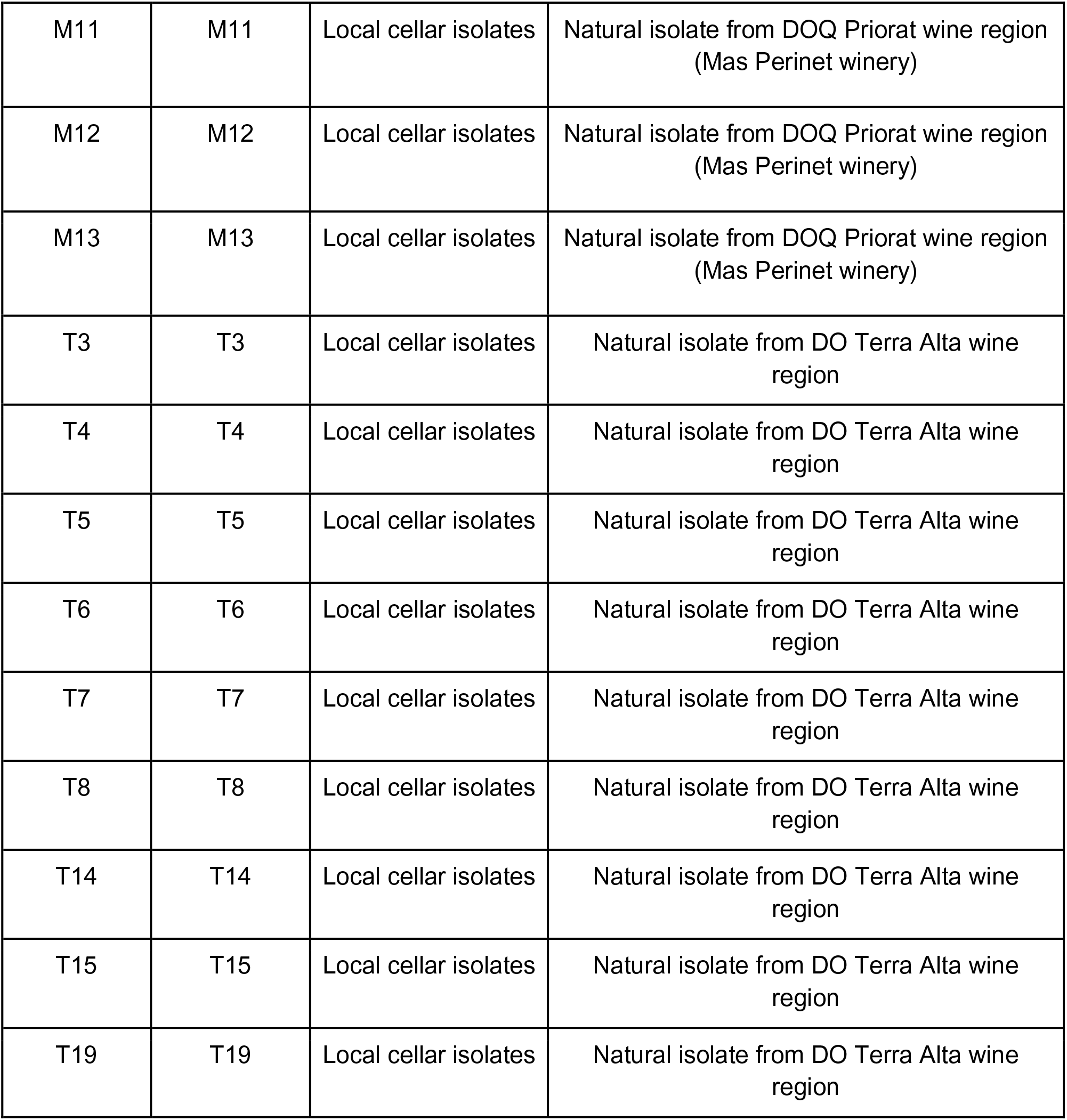
Strains used in the study. Wine yeasts = commercial wine yeasts owned or marketed by Lallemand Inc (Canada). Vineyard yeasts = natural, non-commercialized vineyard yeasts; from grapes or vineyard soil in the Priorat wine-making region in Catalonia, and identified as *S. cerevisiae* using restriction fragment length polymorphisms.

**Table S2.** Selection regimes.

| Name | Background | Description & Rationale |
| --- | --- | --- |
| NCR relaxation | *S. cerevisiae* use of organic nitrogen sources common in grape must (proline, arginine) is often repressed by the presence of inorganic ammonium. Ammonium stimulates the nitrogen catabolite repression system (NCR), which transcriptionally and post-transcriptionally represses genes required for uptake and catabolism of e.g. proline and arginine. Incomplete use of these nitrogen sources leads to stuck fermentations, inability to consume all sugar and invasion of the must by spoiling microorganisms<sup>97</sup>. Amines ($RCH_2NH_2$), such as methylamine, likewise stimulates NCR and represses proline and arginine use. *Saccharomyces cerevisiae* lacks the amine oxidases *AMO1* and *AMO2* and is therefore not capable of using methylated amines as nitrogen sources (PMID 14004755, PMID 721293, 24608175). Methylamine therefore suppresses the use of proline and arginine by activating NCR but does provide the nitrogen that yeast requires for growth. | Select evolving populations on a synthetic grape must medium that only contains arginine and proline as nitrogen sources (15mg/L each) and where use of arginine and proline is heavily repressed by the presence of methylamine (10 g/L). We expect a subset of populations to adapt by reducing the activation of NCR by methylamine, allowing yeast to better use arginine and proline to support growth. Because of the interconnectivity of intracellular nitrogen metabolite pools, we also expect increased flow through the proline and arginine catabolism systems to spill over into higher flows through other nitrogen metabolic pathways, such as those involving branched chain amino acids. This may result in increased production of aroma compounds, and enhanced wine taste and smell. |
| GR relaxation | *S. cerevisiae* often use fructose in grape must less well in the presence of glucose. Incomplete use of fructose often leads to stuck fermentations, inability to consume all sugar and invasion of the must by spoiling microorganisms<sup>98</sup>. 2-deoxy-D-glucose, which has the 2-hydroxyl group replaced by hydrogen, possesses many of the properties of glucose, but because it competitively inhibits the production of glucose-6-phosphate<sup>99</sup>, it cannot undergo glycolysis and cannot be used carbon or energy source by *Saccharomyces cerevisiae*. | Select evolving populations on a synthetic grape must medium that only contains fructose as carbon source and where use of fructose is disfavoured due to the presence of 2-deoxy-D-glucose (2g/L). We expect a subset of populations to adapt by relaxation of glucose repression, allowing earlier and faster use of fructose. Such strains will be particularly useful for re-starting fermentations that have stuck due to incomplete use of fructose. |
| Glutathione production | Glutathione prevents oxidation of wine but is rarely present in notable concentrations in grape must. Yeast produces glutathione from glutamate, cysteine and glycine in a two-step reaction catalyzed by *GSH1* and *GSH2* and can be excreted<sup>100</sup>. The intracellular demand for, and production of glutathione, increases heavily under oxidizing conditions imposed by addition of the oxidizing agent diamide<sup>101</sup>. | Select evolving populations on a synthetic grape must medium only containing glutamate, cysteine and glycine (equal proportions; standard amounts of nitrogen retained 140 mg N/L) as nitrogen sources, increasing their intracellular pools and the flow through glutathione biosynthesis. Diamide is added at growth limiting concentrations (1.5 mM) to simultaneously increase demand for high intracellular glutathione levels. We expect a subset of populations to adapt by mutations that directly (e.g. glutathione biosynthesis) or indirectly (e.g. precursor uptake) increase net production of glutathione. |
| Aroma production | Wine aroma compounds, e.g. fusel alcohols, produced by wine yeasts are bio-synthesized via pathways, e.g. the Ehrlich pathway <sup>102</sup>, originating in the intracellular pools of branched chain amino acids, such as isoleucine, phenylalanine and valine. | Select evolving populations on a synthetic grape must medium that only contains isoleucine, phenylalanine and valine (equal proportions; standard amount of nitrogen retained) as nitrogen sources. We expect a subset of populations to adapt by mutations that directly (e.g. the Ehrlich pathway) or indirectly (e.g. BCAA uptake) increases the production of fusel alcohols. |
| Ethanol tolerance | Due to climate change, the sugar content of grapes creeps upwards. Because, yeast ferments sugar to ethanol, so do the ethanol content of wine. To avoid stuck fermentations, sugar left in the wine, and wine spoilage, this demands wine yeast capable of growth and fermentation at high concentrations of ethanol. The need is particularly pronounced for sparkling wine, which is produced by two serial fermentations of grape must. The second fermentation is initiated by inoculating yeast into wine already high in ethanol content, exposing the yeast to an ethanol shock unless it is pre-adapted (ref). Ethanol is volatile and evaporates under open environment selection experiments. Much less volatile alcohols, such as 1-butanol (bp 118 C vs. 78 C of EtOH), have cellular effects that mimic those of ethanol, and strains resistant to 1-butanol tend to be resistant to ethanol. | Select evolving populations on a synthetic grape must medium supplemented by growth limiting (1.3% v/v) concentrations of 1-butanol. We expect a subset of populations to adapt by increasing their tolerance to alcohols in general, including to ethanol. |
| Sugar tolerance | Due to climate change, the sugar content of grapes creeps upwards <sup>103</sup>. High sugar content exposes yeast to osmotic and ethanol stress and affects sugar uptake by emphasizing the importance of low affinity hexose transporters. | Select evolving populations on a synthetic grape must medium with high (35%; standard proportions of glucose and fructose retained) sugar concentrations. We expect some populations to adapt by increasing their tolerance to osmotic and ethanol stress, and to be able to take up sugar faster and more efficiently at higher sugar concentrations. |
| Vitamin starvation | Vitamins are required for yeast growth and metabolism because of their roles as enzymatic cofactors. In many grape musts, access to vitamins constrains yeast growth and metabolism lead to stuck fermentation, sugar and nitrogen left in the medium, and wine spoilage <sup>97</sup>. The problem is exacerbated by climate change, which leads to faster grape maturation and lower vitamin content of mature grapes (Ref. Moretti). | Select evolving populations on a synthetic grape must medium poor in vitamins (1% of normal; standard proportions retained). We expect some populations to adapt by increasing their capacity to grow and metabolize at low vitamin concentrations. |
| Nitrogen starvation | Grape must is often rich in carbon, but poor in nitrogen, leading to nitrogen starved yeast, stuck fermentations, sugar left in the wine and wine spoilage <sup>104</sup>. | Select evolving populations on a synthetic grape must medium poor in nitrogen (10% of normal; standard proportions retained). We expect some populations to adapt by increasing their capacity to grow and metabolize at low nitrogen concentrations. |

**Table S3.** Media composition for evolution and side-effect environments. Synthetic grape must^32^ was the background media throughout all the experiments.

| Abbreviation | Description | Selection regime | Side-effect |
| --- | --- | --- | --- |
| S-NCR | SGM adjusted as [arginine (15 mg N/l) + proline (15 mg N/l)] as sole N sources + 1% Methyl amine | NCR relaxation |  |
| S-GR | SGM adjusted as 20% Fru. + 2-deoxy glucose (0.2 g/100 ml) | GR relaxation |  |
| S-Gluth | SGM adjusted as [Glycine (47 mg N/l) + Glutamine (47 mg N/l) + Cysteine (47 mg N/l)] as sole N source + 1.5 mM Diamide | Glutathione production |  |
| S-Aroma | SGM adjusted as [Valine (10 mg N/l) + Iso-leucine (10 mg N/l) + Phenylalanine (10 mg N/l)] as sole N sources | Aroma production |  |
| S-But | SGM + 1.3% 1-butanol (v/v) | Ethanol tolerance |  |
| S-35Sug | SGM adjusted as 17.5% Glu. + 17.5% Fru as C source | Sugar tolerance |  |
| S-Vit | SGM adjusted as 1% of normal vitamin concentration | Vitamin starvation |  |
| S-Nit | SGM adjusted as 10% of normal amino acid concentration | Nitrogen starvation |  |
| S-40Sug | SGM adjusted as 20% Glu. + 20% Fru as C source | --- | C utilization |
| S-Fru | SGM adjusted as 20% Fructose as a C source | --- | C utilization |
| S-Lac | SGM adjusted as 2% Lactose as a C source | --- | C utilization |
| S-Mal | SGM adjusted as 2% Maltose as a C source | --- | C utilization |
| S-Man | SGM adjusted as 2% Mannose as a C source | --- | C utilization |
| S-Raf | SGM adjusted as 2% Raffinose as a C source | --- | C utilization |
| S-Gly | SGM adjusted as 2% Glycerol as a C source | --- | C utilization |
| S-Gal | SGM adjusted as 2% Galactose as a C source | --- | C utilization |
| S-Ala | SGM adjusted as Alanine (mg N/l) as sole N source | --- | N utilization |
| S-Asp | SGM adjusted as Aspartic acid (mg N/l) as sole N source | --- | N utilization |
| S-Glt | SGM adjusted as Glutamine (mg N/l) as sole N source | --- | N utilization |
| S-Glyc | SGM adjusted as Glycine (mg N/l) as sole N source | --- | N utilization |
| S-GABA | SGM adjusted as GABA (mg N/l) as sole N source | --- | N utilization |
| S-Met | SGM adjusted as Methionine (mg N/l) as sole N source | --- | N utilization |
| S-Pro | SGM adjusted as Proline (mg N/l) as sole N source | --- | N utilization |
| S-Phe | SGM adjusted as Phenylalanine (mg N/l) as sole N source | --- | N utilization |
| S-Urea | SGM adjusted as Urea (mg N/l) as sole N source | --- | N utilization |
| S-Val | SGM adjusted as Valine (mg N/l) as sole N source | --- | N utilization |

**Table S4.** List of strains used in validations at semi-industrial fermentation scale.

| Founder strain | Evolved population | Selection environment |
| --- | --- | --- |
| M3 | M3 (2,35) | High Sugar |
| E2 | E2 (26,37) | High Sugar |
| G11 | G11 (23,22) | High Sugar |
| M12 | M12 (18,23) | High Sugar |
| E9 | E9 (0,7) | High Ethanol |

**Table S5.** ALE endpoint populations for which glutathione levels was validated.

| Founder strain | ALE population evolved for improved glutathione use |
| --- | --- |
| E6 | E6 (30,7) |
|  | E6 (31,30) |
| G10 | G10 (19,18) |
| T3 | T3 (15,35) |
|  | T3 (15,32) |
| T5 | T5 (11,15) |
| T4 | T4 (0,43) |
|  | T4 (0,19) |
| E11 | E11 (22,31) |
| G9 | G9 (13,1) |
| T15 | T15 (10,19) |
| M3 | M3 (1,35) |
| G6 | G6 (1,36) |
| M4 | M4 (15,28) |
| E7 | E7 (15,46) |
| E5 | E5 (15,39) |
| M7 | M7 (9,24) |
| D12 | D12 (21,34) |

**Table S6.** **Excel file:** 26 fast adapting populations sequenced.

**Table S7.** **Excel file:** point mutations or small insertions/deletions in adapting populations.

**Table S8.** **Excel file:** Loss-of-Heterozygosity (LOH).

